# Gene Editing of Caffeic-O-methyltransferase (COMT1) in the model grass Setaria viridis Improves Biomass Saccharification Without Compromising Plant Growth or Abiotic Stress Tolerance

**DOI:** 10.64898/2025.12.17.694996

**Authors:** Fernanda de Oliveira Menezes, Karoline Estefani Duarte, Gabriel Garon Carvalho, Leonardo Dario Gomez, Igor Cesarino, Wagner Rodrigo de Souza

## Abstract

Second-generation bioethanol production is limited by the recalcitrance of lignocellulosic biomass, largely driven by lignin content and composition. Genetic strategies targeting lignin biosynthesis offer a promising alternative to improve biomass digestibility without relying on energy-intensive pretreatments. Here, we used CRISPR/Cas9 editing to disrupt the caffeic-O-methyltransferase gene (*COMT1*) in *Setaria viridis*, a C₄ monocot model related to major bioenergy crops. *COMT1* was selected based on its strong expression in lignifying tissues and its central role in the biosynthesis of syringyl (S) units, guaiacyl (G) units, and the flavone tricin, a non-canonical lignin monomer in grasses. Edited plants showed no visible growth defects, maintaining normal development and salt-stress tolerance. Chemical analyses revealed a drastic reduction in S units and tricin, accompanied by moderate but significant decreases in G and *p*-hydroxyphenyl (H) monomers. Despite these compositional changes, total soluble lignin content was only slightly reduced, suggesting compensatory incorporation of non-canonical monomers. Metabolome analysis of edited plants suggested a redirection of carbon flux towards phenylpropanoid and flavonoid pathways. These modifications resulted in a substantial increase in saccharification efficiency, demonstrating that *COMT1* disruption can enhance biomass digestibility without compromising plant viability. Our findings highlight *COMT1* as a key target for engineering improved feedstocks for bioenergy production.

**Highlights:**

CRISPR knockout of *COMT1* in *Setaria viridis* markedly enhances biomass digestibility by reducing lignin monomers, without compromising plant growth or salt-stress tolerance.
Genome editing of *COMT1* in *Setaria viridis* reveals unexpected lignin plasticity, reducing S, G, H and tricin units and improving saccharification efficiency without developmental penalties.
Combined thiacidolysis/GC-MS and metabolomics suggests a compensatory mechanism in the lignin portion of the *S. viridis* cell wall.

## 1. Background

As highlighted in the *World Energy Outlook* (U.S. Energy Information Administration, 2021), the energy sector accounts for nearly three-quarters of all anthropogenic greenhouse gas (GHG) emissions worldwide. This alarming statistic underscores the urgent need for transformative changes in how energy is produced and consumed to mitigate climate change and promote long-term ecological resilience effectively. Within this context, the development of sustainable energy systems has gained prominence as a strategic pathway to curb emissions. Among the proposed alternatives, second-generation (E2G) bioethanol derived from lignocellulosic biomass such as agricultural residues has attracted significant international attention as a renewable, low-emission substitute for fossil fuels (Reshmy *et al*., 2023; Yılbaşı, 2025).

E2G is produced from lignocellulosic biomass derived mainly from agricultural residues such as sugarcane bagasse. The production process begins with a physical and/or chemical pretreatment aimed at reducing biomass recalcitrance by disrupting the lignocellulosic matrix. This step is essential to enhance biomass digestibility, as the rigid cell wall structure limits enzymatic accessibility. Following pretreatment, enzymatic hydrolysis converts structural carbohydrates into fermentable sugars, which are subsequently fermented to ethanol. Improving biomass digestibility remains a major technological bottleneck in E2G production, with a direct impact on process efficiency and economic viability (Liu *et al*., 2019, Preprint; Kordala *et al*., 2024, Preprint).

Lignocellulosic biomass primarily comprises three biopolymers that form plant cell walls: cellulose, hemicelluloses, and lignin (Ma *et al*., 2019, Preprint; Shi *et al*., 2019). The cellulose polymer forms a scaffold of microfibrils embedded in a matrix of hemicelluloses, which are interlinked with lignin. Lignin, a complex phenolic polymer mainly composed of *p*-hydroxyphenyl (H), guaiacyl (G), and syringyl (S) units, confers mechanical strength to the cell wall, facilitates water transport, and plays key roles in plant defense mechanisms against biotic and abiotic stressors. However, from a bioindustrial perspective, lignin represents a major barrier to biomass deconstruction. Its hydrophobic nature and cross-linking capacity restrict enzymatic access to polysaccharides, thereby reducing sugar release during hydrolysis and compromising fermentation efficiency (Broda *et al*., 2022, Preprint; Abolore *et al*., 2024, Preprint).

Although physicochemical pretreatments can partially mitigate biomass recalcitrance, they often require high energy input, increase processing costs, and may produce environmentally harmful byproducts (Basak *et al*., 2023; Abolore *et al*., 2024, Preprint). As an alternative, targeted genetic modifications of cell wall components have emerged as a promising strategy to enhance biomass digestibility (Damm *et al*., 2016, Preprint; Rao and Dixon, 2018, Preprint). Reducing lignin content or altering its composition enhances the release of fermentable sugars from structural carbohydrates, representing a sustainable approach to improving biomass conversion efficiency and reducing second-generation bioethanol production costs (Mosier *et al*., 2005; Tye *et al*., 2016, Preprint). However, these genetic modifications often lead to growth penalties, such as reduced biomass yield and increased susceptibility to abiotic stresses, which limit their scalability. Therefore, optimizing the balance between lignin modification, plant productivity, and stress resilience is essential for advancing the next generation of bioenergy crops (De Vries *et al*., 2018).

Currently, twelve enzyme families have been identified as key players in the biosynthetic pathway towards canonical lignin monomers (Figure 1). The primary precursor phenylalanine is deaminated into cinnamic acid, which is converted into distinct hydroxycinnamyl alcohols (generally known as monolignols) via hydroxylations/methylations of the aromatic ring and reduction of the propane tail. Multiple routes have been described to contribute to the biosynthesis of monolignols in different plant species, including a tyrosine by-pass via tyrosine ammonia-lyase activity (Barros *et al*., 2016), the “acid pathway” via bifunctional ascorbate peroxidase activity (Barros *et al*., 2019), and the highly evolutionarily conserved “shikimate shunt” (Dixon and Barros, 2019, Preprint). The complexity of lignin biosynthesis drastically increases when considering lignin monomers beyond the canonical monolignols, as 49 metabolites have been detected to be incorporated into lignin polymers in different cellular contexts and plant species (Pesquet *et al*., 2025, Preprint). Among key lignin biosynthetic enzymes, Caffeic Acid O-Methyltransferase (COMT, E.C. 2.1.1.68) is a *S*-adenosyl-L-methionine-dependent *O*-methyltransferase involved in multiple steps of the pathway. Originally, COMT was thought to catalyze the methylation of caffeic and 5-hydroxyferulic acid, but latter studies demonstrated that this enzyme preferentially methylate 5-hydroxyconiferaldehyde and/or 5-hydroxyconiferyl alcohol to produce sinapaldehyde and/or sinapyl alcohol, respectively (Humphreys J.M and Chapple C, 2002). Thus, together with Ferulic Acid 5′-Hydroxylase (F5H), COMT constitutes the specific pathway towards the biosynthesis of syringyl lignin units. More recently, COMT-mediated methylation of caffeic acid into ferulic acid was demonstrated to be of great importance for lignification in grasses, being proposed as the most favored route towards G and S units in this group of plants (Barros *et al*., 2019). Additionally, COMT was demonstrated to play dual roles in the lignin pathway in grasses, as it catalyzes not only the methylation steps within the S-lignin branch but also the methylation steps towards the biosynthesis of the flavone tricin (Eudes *et al*., 2017), an authentic lignin monomer in grasses (Lan *et al*., 2015). This enzyme also plays an important role in responses to abiotic stresses, such as drought and salinity, through the synthesis of melatonin. COMT catalyzes the conversion of N-acetyl-5-hydroxytryptamine into melatonin, an important regulator of oxidative stress and redox balance in plants, helping to protect cells against damage caused by abiotic stress (Li *et al*., 2022). Due to its key position in the lignin biosynthetic pathway, the *COMT* gene is considered a suitable target for regulating lignin monomers aimed at reducing biomass recalcitrance.

**Figure 1.**
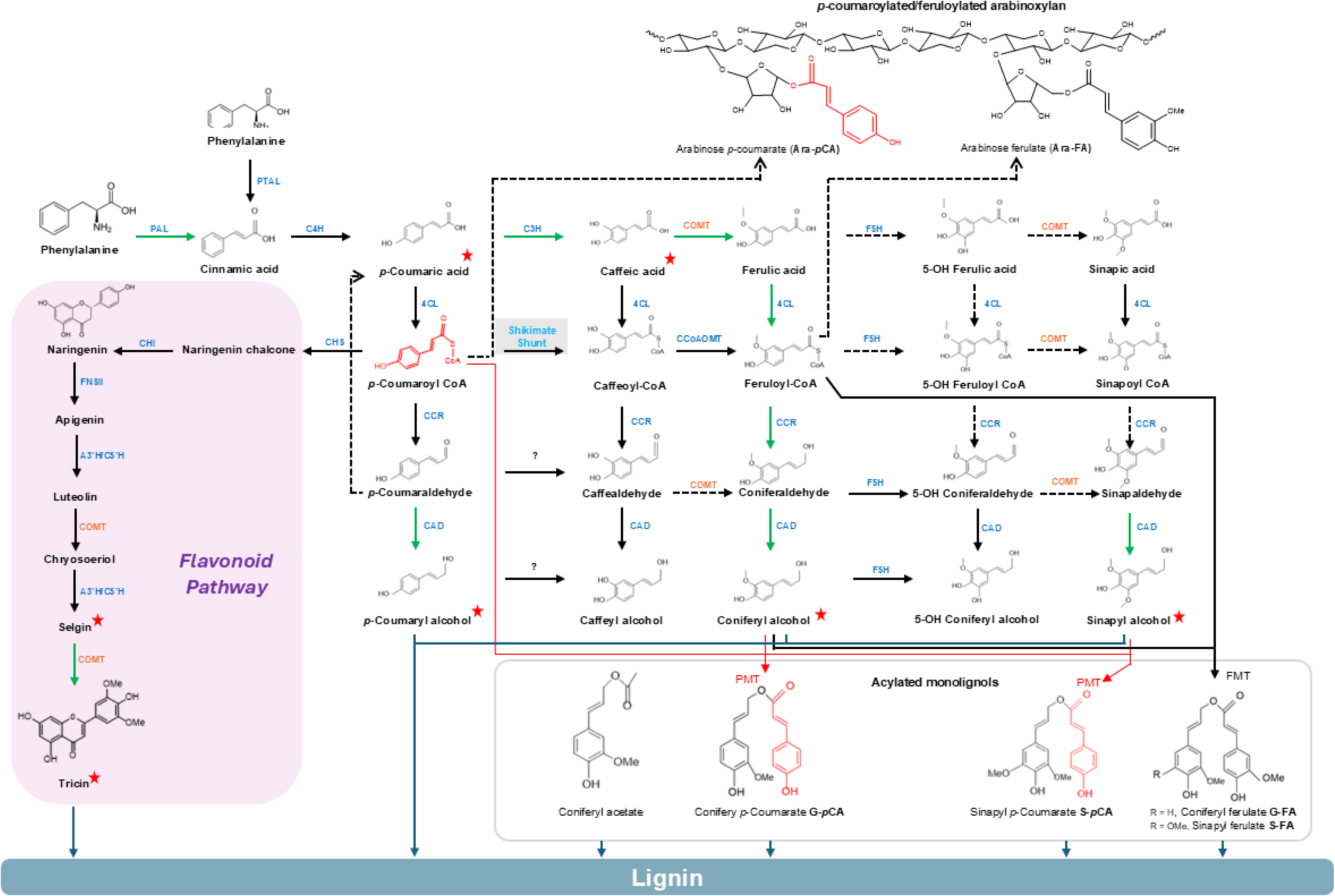
Lignin biosynthetic pathway in grasses. Current model of lignin biosynthesis encompassing the phenylpropanoid and flavonoid pathways. Preferential carbon flux is indicated by green arrows, while dashed arrows denote unconfirmed steps. The “shikimate shunt” of the phenylpropanoid pathway is highlighted in gray. Solid arrows indicate experimentally confirmed enzymatic steps, and dashed arrows represent suggested conversions. **Phenylpropanoid-related enzymes:** PAL (phenylalanine ammonia-lyase), PTAL (phenylalanine/tyrosine ammonia-lyase), C4H (cinnamate 4-hydroxylase), 4CL (4-hydroxycinnamate:CoA ligase), C3H (4-coumarate 3-hydroxylase/ascorbate peroxidase), CSE (caffeoyl shikimate esterase), HCT (hydroxycinnamoyl-CoA:shikimate/quinate hydroxycinnamoyl transferase), C3’H (4-coumaroyl shikimate 3’-hydroxylase), CCoAOMT (caffeoyl-CoA 3-O-methyltransferase), CCR (cinnamoyl-CoA reductase), F5H (ferulate/coniferaldehyde 5-hydroxylase), COMT (caffeic acid O-methyltransferase), CAD (cinnamyl alcohol dehydrogenase), PMT (p-coumaroyl-CoA:monolignol transferase), FMT (feruloyl-CoA:monolignol transferase). **Flavonoid- and flavone-related enzymes:** CHS (chalcone synthase), CHI (chalcone isomerase), FNSII (flavone synthase II), A3’H/C5’H (apigenin 3’-hydroxylase/chrysoeriol 5’-hydroxylase). The red star marks the intermediates that were altered in *COMT1*-edited plants.

Studies involving the negative regulation of the *COMT* gene using RNA interference (*RNAi*) techniques in plants of *Brachypodium distachyon* (Ho-Yue-Kuang *et al*., 2016), *Panicum virgatum* (Wu et al., 2018), and *Hordeum vulgare* (Daly *et al*., 2019) observed significant changes in lignin composition and biomass digestibility. However, the negative regulation of *COMT* in *Zea mays* and *Sorghum bicolor* plants resulted in negative effects on biomass yield (Miller *et al*., 1983; Sattler *et al*., 2010, Preprint). In sugarcane (*Saccharum* sp.), Jung et al. (2012) silenced the COMT gene, addressing biomass digestibility. Although these plants showed increased biomass saccharification, there was also a decrease in productivity. In a more recent study, Kannan et al. (2018) used TALEN-mediated mutagenesis for the knockout of 100 copies/alleles of the *COMT* gene in sugarcane increasing saccharification without compromising biomass.

*Setaria viridis* is an emerging monocot model plant for molecular and genetic research. This short, fast-growing, C₄ species has a fully sequenced genome (Bennetzen *et al*., 2012). It belongs to the *Panicoideae* subfamily, which includes the bioenergy crops *Sorghum bicolor, Zea mays, and Saccharum sp.* In a study conducted by Ferreira et al. (2019), six *COMT* genes were identified in the *Setaria viridis* genome. According to this study, the *COMT1 and COMT6* genes showed phylogenetic proximity to the *COMT* genes involved in lignification in *Brachypodium distachyon, Zea mays, Oryza sativa,* and *Sorghum bicolor* (Piquemal *et al*., 2002*a*; Saballos *et al*., 2008; Koshiba *et al*., 2013; Trabucco *et al*., 2013*a*; Ho-Yue-Kuang *et al*., 2016). Furthermore, it was observed that most of the *COMT* genes showed almost null expression in the elongated internode of *Setaria viridis,* except for *COMT1,* which demonstrated strong and preferential expression in the transition and maturation zones Ferreira et al. (2019) This work shows that *COMT1* is the main candidate gene to play a crucial role in lignification during development in *Setaria viridis*.

Here, we demonstrate that CRISPR/Cas9 knockout of the *COMT1* gene in *Setaria viridis* represents a promising strategy to enhance biomass digestibility. The targeted genome editing significantly improved saccharification efficiency without compromising overall plant viability. Notably, despite substantial alterations in lignin composition, the mutant plants maintained normal growth and did not exhibit increased sensitivity to salt stress. These findings underscore the feasibility of engineering lignin pathways without detrimental effects on plant development.

## 2. Materials and methods

### 2.1. Strategies for *COMT1* gene editing in *Setaria viridis* using CRISPR/Cas9

The sequence of the target gene, *COMT1* from *Setaria viridis*, was retrieved from the public database Phytozome v13 (https://phytozome-next.jgi.doe.gov/). The genome version used was *Setaria viridis* v4.1, with the corresponding *COMT1* gene ID Sevir. 6G053300. CRISPRPLANT v.2 software (Minkenberg *et al*., 2019) was utilized to identify the sgRNAs corresponding to the selected gene-editing target, focusing on the region immediately upstream of the PAM sequence (5′-NGG), specific to each gene region to be deleted. The software also enabled the analysis of off-target regions for the selected sgRNAs, and target sites with low probabilities of mis-targeting were chosen for gene editing. The sequences of the sgRNAs for *COMT1* are available in Supplementary Table 1.

After identifying the sgRNAs, we proceeded with the construction of vectors for the genetic transformation of *S. viridis*. The selected sequences were cloned into a CRISPR vector (E856; p6i2xoR-UcasW-U6Os4), with codon optimization for monocot species. Cloning was performed by DNA Cloning Service (Hamburg, Germany). The schematic representation of the optimized CRISPR/Cas9 vector is shown in Supplementary Figure 1A, and a schematic representation of the sgRNA target sites in the *COMT1* gene is shown in Supplementary Figure 1B.

### 2.2. Genetic transformation of *Setaria viridis* and CRISPR/Cas screening

The CRISPR vector was introduced into *Agrobacterium tumefaciens* (strain EAH105α) via heat shock, and genetic transformation of *S. viridis* A10.1 was carried out by co-inoculating embryogenic callus with *A. tumefaciens*, following the protocol adapted from (Martins *et al*., 2015). Transformed plants were confirmed by PCR using primers specific to the amplification of the *hptII* (244 bp) and *Cas9* (500 bp) genes. Primers used to confirm the transformed events are listed in Supplementary Table 2. To detect insertions or deletions (indels) in the target sites, PCR amplification was performed to generate gene fragments spanning DSB sites (gRNA regions), followed by PCR product purification and Sanger sequencing to validate potential mutations. Chromatograms from sequencing were visualized using BioEdit® software (version 7.2.5). For mutation analysis, we utilized SYNTHEGO software, specifically the ICE (Inference of CRISPR Edits) Analysis tool, as described by (Hsiau *et al*., 2018, Preprint) (available at https://ice.synthego.com/#/). Samples with R² values below 0.8 were excluded due to the lower reliability of the results. In addition to R², we also considered the percentages of indels formed, the nature of these indels, and the modifications induced by mutations in the chromatogram. Thus, our analysis included both the percentage of indels and the knockout score.

### 2.3. Plant material and growth conditions

Independent lines with frameshift mutations were propagated up to the T_2_ generation. They were cultivated in a substrate mixed with vermiculite at a 3:1 ratio, in a controlled environment at 28 ± 2 °C, with a photoperiod of 16 hours of light and 8 hours of darkness. Irrigation was performed six times a week, maintaining field capacity, while a nutrient solution containing 1 mL of Plant Magic Plus Bio Silicon® per liter of water was applied weekly. The fertilizer composition included 11.7% potassium oxide, 14.3% water-soluble silicon, and 0.05% water-soluble EDTA-chelated iron. Cell wall characterization was conducted with sampling during the senescence phase, when the plants were naturally aging. Plant material (leaves and stems) was processed into powder using a ball mill. Each mutant was treated as an individual biological replicate, and four technical replicates were performed for subsequent analyses

### 2.4. Cell wall characterization

Independent lines with frameshift mutations were propagated up to the T_2_ generation. They were cultivated in a substrate mixed with vermiculite at a 3:1 ratio, in a controlled environment at 28 ± 2 °C, with a photoperiod of 16 hours of light and 8 hours of darkness. Irrigation was performed six times a week, maintaining field capacity, while a nutrient solution containing 1 mL of Plant Magic Plus Bio Silicon® per liter of water was applied weekly. The fertilizer composition included 11.7% potassium oxide, 14.3% water-soluble silicon, and 0.05% water-soluble EDTA-chelated iron. Cell wall characterization was conducted with sampling during the senescence phase, when the plants were naturally aging. Plant material (leaves and stems) was processed into powder using a ball mill. Each mutant was treated as an individual biological replicate, and four technical replicates were performed for subsequent analyses.

### 2.5. Preparation of alcohol insoluble residue (AIR)

Alcohol-insoluble residue (AIR) was prepared as described by Möller et al. (2022), by adding 2 mL of water to 100 mg of powdered plant material in a 2 mL Eppendorf tube. The powder was resuspended in the water and shaken by inverting the tube for 30 seconds. The plant material was then pelleted by centrifugation at 4000 *g* for 10 minutes. The supernatant was removed and replaced with 2 mL of 96% ethanol, followed by two additional ethanol washes. The resulting residue was allowed to dry under ambient conditions.

### 2.6. Lignin quantification by acetyl bromide method

The determination of soluble lignin was carried out by the acetyl bromide method, as described by Liu et al. (2016). Precisely 5 mg of AIR was added to a centrifuge tube with a screw cap, along with 250 µL of acetyl bromide solution (25% acetyl bromide and 75% glacial acetic acid, v/v). The mixture was heated at 50 °C for three hours. After cooling the samples to room temperature, the resulting liquid was transferred to a 5 mL volumetric flask. Next, 1 mL of 2 M NaOH and 175 μL of 0.5 M hydroxylamine hydrochloride (HCl) were added to the reaction. The final volume of the reaction was adjusted precisely to 5 mL using glacial acetic acid. The absorbance of the supernatant was measured on a spectrophotometer at 280 nm. The calculation of the total lignin content in the dry sample was performed using the following formula, where the coefficient used in the calculation was 17.75, specific for grasses (Foster *et al*., 2010; Dwivedi *et al*., 2024).

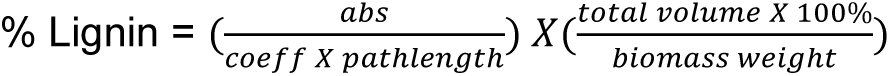

### 2.7. Analysis of lignin composition by thioacidolysis method

Lignin composition analysis was also performed using the method described by Chen et al. (2021), with some modifications. Briefly, 1 mL of thioacidolysis reagent (composed of 100 mL of 1,4-dioxane, 6.25 mL of boron trifluoride diethyl ether, and 25 mL of ethanethiol, in a v/v ratio) was added to 2.5 mg of AIR. 4,4-Ethylidenebisphenol was added as an internal standard at a final concentration of 12 µg/mL. The reaction was incubated for 4 hours at 100 °C. After cooling, 400 µL of the solution was transferred to glass vials, and 190 µL of 0.4 M NaHCO_3_ were added. The reaction mixture was incubated overnight at 55 °C. The resulting pellet was then homogenized in 100 µL of derivatization reagent (1:1 pyridine and BSTFA with 1% TCM v/v) at 55 °C for 30 minutes prior to GC-MS analysis. For tricin analysis, 400 µL of the thioacidolysis reaction was transferred to 1.5 mL tubes, followed by the addition of 190 µL of 0.4 M NaHCO3. This mixture was incubated overnight at 55 °C. The resulting pellet was homogenized in a 1:1 (v/v) methanol:DMSO solution. Samples were sonicated for 10 seconds, centrifuged at 22,000 *g* for 10 minutes, and 100 µL of the supernatant was transferred to vials for HPLC analysis. An Agilent Zorbax Eclipse Plus C18 column (2.1 mm × 50 mm, 1.8 µm) was used to separate the compounds. The mobile phases used were: (A) 0.1% formic acid in water (v/v) and (B) 0.1% formic acid in acetonitrile (v/v). The column thermostat was maintained at 30 °C with a solvent flow rate of 0.45 mL/min. The elution gradients were as follows: initially 6% (B) for three minutes, followed by an increase from 6% to 95% (B) over 5 minutes. The mobile phase was maintained at 95% (B) for two minutes, followed by a return from 95% to 6% (B) in one minute, and a final phase of two minutes at 6% (B). The total run time was 13 minutes.

### 2.8. Determination of total *p*-CA and FA by alkaline hydrolysis

The extraction of *p-*coumaric acid (*p*-CA) and ferulic acid (FA) followed the protocol established by Möller et al. (2022). Approximately 10 mg of AIR were incubated with 1M NaOH at 30 °C for 12 hours. The reaction was then neutralized with 99% trifluoroacetic acid (TFA) and subjected to a liquid-liquid extraction using 700 μL of butanol. After the evaporation of butanol, the resulting residue was resuspended in a 50% methanol solution and analyzed by high-performance liquid chromatography (HPLC) with UV detection. The injection volume was 10 μL. Component separation was performed using an XBridge BEH Shield RP18 chromatographic column (Waters, 186003149) with a porosity of 130 Å, a particle size of 3.5 μm, and dimensions of 1 mm × 100 mm. The flow rate was set to 1.23 mL/min, and a binary gradient of 0.1% acetic acid and methanol was applied. The gradient program increased the methanol concentration from 5% to 70% over 0–30 minutes, then from 70% to 90% between 30 and 35 minutes and finally returned it to 5% between 35 and 40 minutes. Standard curves were generated using commercial *p*-CA and FA standards (Sigma) over a concentration range of 0.02–3.28 µg/mL for *p*-CA and 0.02–3.88 µg/mL for FA.

### 2.9. Determination of *p*-CA-arabinose and FA-arabinose by mild acidolysis

The quantification of *p*-coumaric acid and ferulic acid bound to arabinose, *p*-CA-ara and Fa-ara, respectively, was carried out following the protocol described by Lapierre et al. (2018). Briefly, 1 mL of the acidolysis reagent (composed of dioxane, methanol, and 2M aqueous hydrochloric acid in a 60:30:10 (v/v/v) ratio) was added to 15 mg of AIR in a screw-cap vial. The incubation was carried out for three hours at 80 °C with shaking at 163 *rpm*. After cooling, the reaction was stopped by the addition of 3 mL of water. The resulting diluted mixture was subjected to two extractions with 1 mL of ethyl acetate, followed by the evaporation of the ethyl acetate and dissolution of the pellet in 100 μL of methanol. The analysis was performed by HPLC, using a binary gradient of 0.1% acetic acid and methanol, with an injection volume of 10 μL. Separation of the sample components was achieved using an XBridge BEH Shield RP18 chromatographic column (Waters - 186003149). The gradient was programmed as follows: for the first 0-60 minutes, the methanol concentration was increased from 5% to 65%; from 60-65 minutes, the methanol concentration was increased from 65% to 90%; and from 65-70 minutes, the methanol concentration was returned to 5%.

### 2.10. Monosaccharide analysis

The analysis of the monosaccharide composition of the AIR fraction was performed through the hydrolysis of polysaccharides into monosaccharides. For this, 4 mg of AIR were hydrolyzed in 0.5 mL of 2 M trifluoroacetic acid (TFA) for 4 hours at 100 °C. After cooling to room temperature, the solution was completely evaporated using a centrifugal evaporator. The resulting pellet was washed twice with 2-propanol and evaporated. Subsequently, the material was resuspended in 200 μL of distilled water, and the suspension was filtered through a 0.45 μm PTFE filter. The analysis was carried out by HPAEC using a Dionex Carbopac PA-10 column with pulsed amperometric detection, as described by Möller et al. (2022) and Wannitikul et al. (2024). The quantification of the isolated monosaccharides was performed using an external calibration curve containing nine monosaccharide standards at concentrations of 250, 500, and 700 μM (glucose, fucose, arabinose, galactose, mannose, rhamnose, xylose, galacturonic acid, glucuronic acid).

### 2.11. Determination of cellulose content

The quantification of cellulose was carried out according to the methodology outlined by Acevedo et al. (2019). The TFA pellets, obtained as detailed in the previous section, were subjected to a washing procedure with 1.5 mL of water, followed by three additional washes with 1.5 mL of acetone. Subsequently, the samples were air-dried at room temperature overnight. For complete hydrolysis, 90 µL of 72% (w/v) sulfuric acid were added to the dry pellets. Then, 1.89 mL of water was added, and the mixture was heated at 120 °C for 4 hours. The glucose content in the resulting supernatant was determined using the Anthrone colorimetric assay (Morse, 1947), with a glucose standard curve for quantification. The analysis was performed at a final volume of 1.2 mL, using 40 μL of sample and 800 μL of Anthrone (2 mg/mL). The blue color developed after incubation at 80 °C for 30 minutes. Standard reactions with glucose concentrations ranging from 0.5 μg/mL to 10 μg/mL were included for calibration purposes. The absorbance of the reaction was measured at 620 nm.

### 2.12. Biomass enzymatic hydrolysis (saccharification)

Enzymatic saccharification was carried out according to the procedures outlined by (Gomez *et al*., 2010). Two types of biomasses were used in the saccharification process: untreated biomass and pretreated biomass. Biomass consisted of whole aerial parts of the plants (leaves + stem). For the pretreatment, 0.5 mL of 500 mM NaOH was added to 100 mg of biomass, and the reaction was heated to 90 °C for 30 minutes with occasional stirring. The material was then decanted to the bottom of the tube. Subsequently, it was washed three times with 1 mL of 25 mM sodium acetate buffer at pH 4.5. For enzymatic saccharification, an enzyme cocktail with a 4:1 ratio of Celluclast to Novozyme 188 was used, where Novozyme 188 acted as a cellulase and was derived from *Aspergillus niger* (both enzymes were obtained from Novozymes, Bagsvaerd, Denmark). Before use, the enzymes were filtered using a Hi-Trap desalting column (GE Healthcare, Little Chalfont, Buckinghamshire, UK). The hydrolysis process was carried out in a 25 mM sodium acetate buffer (pH 4.5) at 50 °C for specified time intervals. The determination of sugars released after hydrolysis was performed using a modified version of the method introduced by (Anthon and Barrett, 2002), incorporating 3-methyl-2-benzothiazolinone hydrazone (MBTH). The analysis was conducted in a final volume of 1 mL, with 300 μL of the sample used for the assay. The detection reaction consisted of 0.25 N NaOH, 3 mg/mL MBTH, and 1 mg/mL dithiothreitol (DTT). Color development occurred after incubation at 70 °C for 20 minutes, achieved by adding the oxidizing reagent (0.2% FeNH4(SO4)2, 0.2% sulfamic acid, and 0.1% HCl). Standard reactions with glucose concentrations of 10, 20, 50, and 100 nmol were included for calibration purposes. The absorbance of the reaction was measured at 620 nm. This method was validated to detect a variety of sugars released from the cell wall and demonstrated sensitive detection of several monosaccharides (Gomez *et al*., 2010).

### 2.13. Untargeted metabolome profiling

For metabolome analysis, frozen aerial parts were milled with a Grindomix GM200 using precooled stainless-steel jars (Retsch). Metabolites were extracted using 5% acetonitrile, adding a necessary volume to obtain a concentration of 1 mg/mL. The samples were homogenized in vortex for 10 min. at 6720 x *g*. The supernatant was collected and filtered in microfilter PTFE 0.22 µm. The filtered samples were injected (5 μL) on a UHPLC system (Waters ACQUITY UPLC) equipped with a HSS T3 column (100 mm by 2.1 mm, 1.8 μm; Waters) and coupled to a Waters Synapt XS qToF (quadrupole time-of-flight) hybrid (Waters) mass spectrometer. A gradient of two buffers (A and B) was utilized: buffer A (0.1 % :formic acid, v/v) and buffer B (99:1:0.1 acetonitrile:water:formic acid, pH 3, v/v), as follows: 95% A, 5% B for 3 min, decreased to 50% A and 50% B in 15 min, decreased to 100% B in 18 min and 95% A, 5% B in 20 min, flow rate was 0.280 mL per min. LockSpray ion source was used in negative electrospray ionization mode under the following specific conditions: capillary voltage, 2.50 kV; source temperature, 120°C; desolvation gas temperature, 500°C; desolvation gas flow, 1000 L per h; and cone gas flow, 100 L/h. The mass range was set from 50 to 1200 Da. For DDA-MSMS, the low mass ramp was ramped between 15 and 50 eV. The peak selection was performed when a peak was > 1% of baseline peak spectra. Data processing was performed with Progenesis QI software version 2.4 (Waters). All peaks (mass-to-charge ratio [m/z] features) were integrated in the chromatograms of WT and *SvCOMT* Ev1, Ev2 and Ev3 plants. The following filters were applied to analyze the data: average ion intensity within at least one group >100, P value Student’s t-test < 0.001, and fold change >2 or <0.5 compared to the respective control (WT). Peaks that appeared in the blank (30 blank samples) were also excluded from analysis.

### 2.14. Salt stress assay in hydroponic system

Initially, the seeds were germinated on Petri dishes containing Murashige and Skoog medium with the pH adjusted to 5.8. After germination, the plants were transferred to a hydroponic system and maintained in Hoagland medium (Phytotech Labs, Product ID H353) for 20 days, until the stage just before flowering onset. After the initial growth period, the salinity stress experiment was initiated with three treatments: 0 mM (control), 100 mM, and 250 mM NaCl. The pH of the medium was maintained at 5.8, and the treatment solutions were adjusted according to the specified NaCl concentrations. The growth environment was controlled to maintain a photoperiod of 16 hours of light and 8 hours of darkness, with a constant temperature of 26 °C during both day and night. The nutrient solutions were renewed every 4 days to ensure the maintenance of the desired nutrient and salt concentrations. Plants were collected in quadruplicate after 5 days and 20 days of treatment exposure.

### 2.15. Determination of relative water content and dry biomass

The relative water content (RWC) of control and stressed plants was determined with 0.2 g of fresh leaves (fresh weight; FW) that were immediately rehydrated in distilled water for 24 h in the dark, when the turgid weight (TW) was recorded. Next, leaves were transferred to an oven set at 60 °C and kept until reaching a constant dry weight (DW). Leaf relative water content was calculated using the following equation (Menezes *et al*., 2020). %RWC=100×(FW−DW)/(TW−DW)

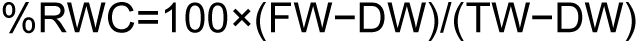

In addition to RWC, the dry biomass of the plants was evaluated after harvesting. Initially, 0.2 g of dry biomass from the aerial part was weighed and then dried in an oven at 60 °C for 24 hours. After this period, the samples were reweighed to determine the final weight.

## 3. Results

3.1. CRISPR/Cas9 of *COMT1* editing resulted in plants with different frameshift mutations

The strategy for *COMT1* gene knockout was based on the non-homologous end joining (NHEJ) DNA repair pathway, which repairs double-stranded DNA breaks (DSBs) generated by nuclease activity through error-prone ligation. Four sgRNAs were designed to direct the CRISPR/Cas9 system to the *COMT1* gene in *Setaria viridis*. The first three (sgRNA1, sgRNA2, and sgRNA3) targeted nucleotides 653–752 within exon 1, a critical coding region essential for gene function, whereas sgRNA4 was designed to target position 1771 in exon 2. This multi-site targeting strategy was intended to maximize knockout efficiency by inducing DSBs at key functional regions of the gene (Supplementary Figure 1B).

*Agrobacterium*-mediated transformation generated multiple independent events, confirmed by PCR and Sanger sequencing. Three independent T₂ lines (Ev.1, Ev.2, and Ev.3) carrying frameshift mutations induced by sgRNAs 1, 2, and 3 were selected for further analysis (Supplementary Figure 2 to 4 and Supplementary Table 3). The selected lines exhibited knockout-scores above 70%, as determined by Synthego ICE analysis. This score represents the proportion of indels predicted to cause functional knockouts—specifically frameshift mutations, indels ≥21 nucleotides, or edits likely to disrupt protein function. Unlike total indel frequency, which includes all insertions and deletions differing from the wild-type sequence, the knockout-score reflects only those mutations expected to abolish gene function. Scores above 50% are generally associated with biallelic mutations (Synthego, 2018; Conant et al., 2022). Based on these criteria, we selected three homozygous lines with distinct knockout-scores (Ev.1, 91%; Ev.2, 75%; and Ev.3, 88%) for subsequent biochemical and phenotypic characterization.

### 3.2. *COMT1* mutants lack the brown midrib phenotype and demonstrate normal growth and development

Reduced lignin content and/or altered lignin composition has been frequently associated with the so-called “brown midrib” phenotype in grasses, a characteristic reddish-brown coloration of leaf blade midribs (Sattler *et al*., 2010, Preprint). This phenotype has been classically applied to identify and isolate lignin mutants in grasses, given that all mutants arising by either spontaneous or chemical mutagenesis were shown to be associated with loss-of-function of lignin-related genes (Sattler *et al*., 2010, Preprint; Tang *et al*., 2014; Li *et al*., 2015; Tetreault *et al*., 2021). After generating knockout mutants for *COMT1* in *S. viridis*, we first assessed whether these plants presented the expected brown midrib phenotype. However, the *COMT1* mutants did not exhibit any obvious visible phenotypes. These plants did not show the presence of a brown central vein in the leaves (Figure 2A) nor significant changes in growth patterns (Figure 2B). After senescence, the mutant lines (Ev.1, Ev.2 and Ev.3) were harvested as described in Section 4. Subsequently, we analyzed the effects of the *COMT1* knockout on the cell wall and biomass saccharification of these plants. Since the individual plants exhibited different editing patterns (Supplementary Figures 2, 3 and 4), each mutant was treated as an independent biological replicate, with four technical replicates in the subsequent experiments.

**Figure 2.**
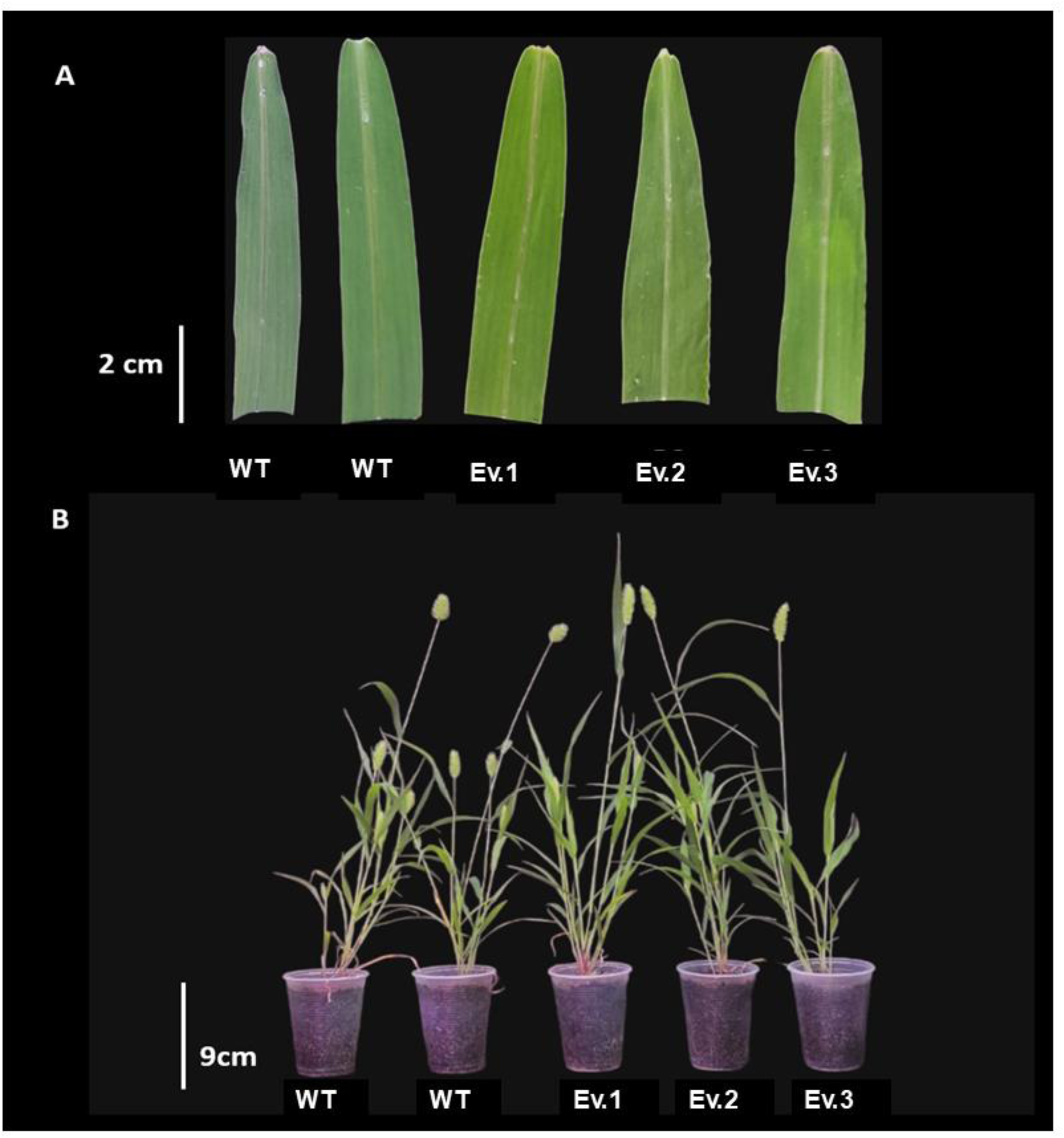
Phenotypes of wild-type (WT) plants and *COMT1* knockout lines. (A) Leaves of WT plants and *COMT1* knockout lines (Ev.1, Ev.2, and Ev.3). The typical symptoms of *bmr* (brown midrib) mutations, characterized by brown coloration in the central vein, were not observed in the knockout lines. (B) Representative image showing no visible growth differences between WT plants and *COMT1* knockout lines (Ev.1, Ev.2, and Ev.3).

### 3.3. Quantification of lignin content and analysis of its monomeric composition reveal differences in *COMT*-edited plants

To investigate the effects of the *COMT1* gene knockout on lignin content and composition, we first quantified the total soluble lignin content in the mutant lines and wild-type plants, following the method described by Liu et al. (2016). A small but statistically significant reduction in total soluble lignin content was observed in the mutant plants, with a decrease ranging from 1.9% to 3.1% compared to the wild-type plants (Figure 3A). The lignin composition was also analyzed using the thioacidolysis method, as described by Chen et al. (2021). The results of this analysis reveal a significant reduction in the G, S, and H monomers in the edited plants compared to the wild-type (WT) plants. For the G monomer, the reduction ranged from 13.3% to 28.8% (Figure 3B). In the case of the S monomer, reductions were more pronounced, ranging from 81.9% to 76.6% (Figure 3C), while the H monomer presented a reduction between 44.1% and 52.5% (Figure 3D). Additionally, tricin, another important component of lignin in grasses, decreased by 44.1% to 51% (Figure 3E).

**Figure 3.**
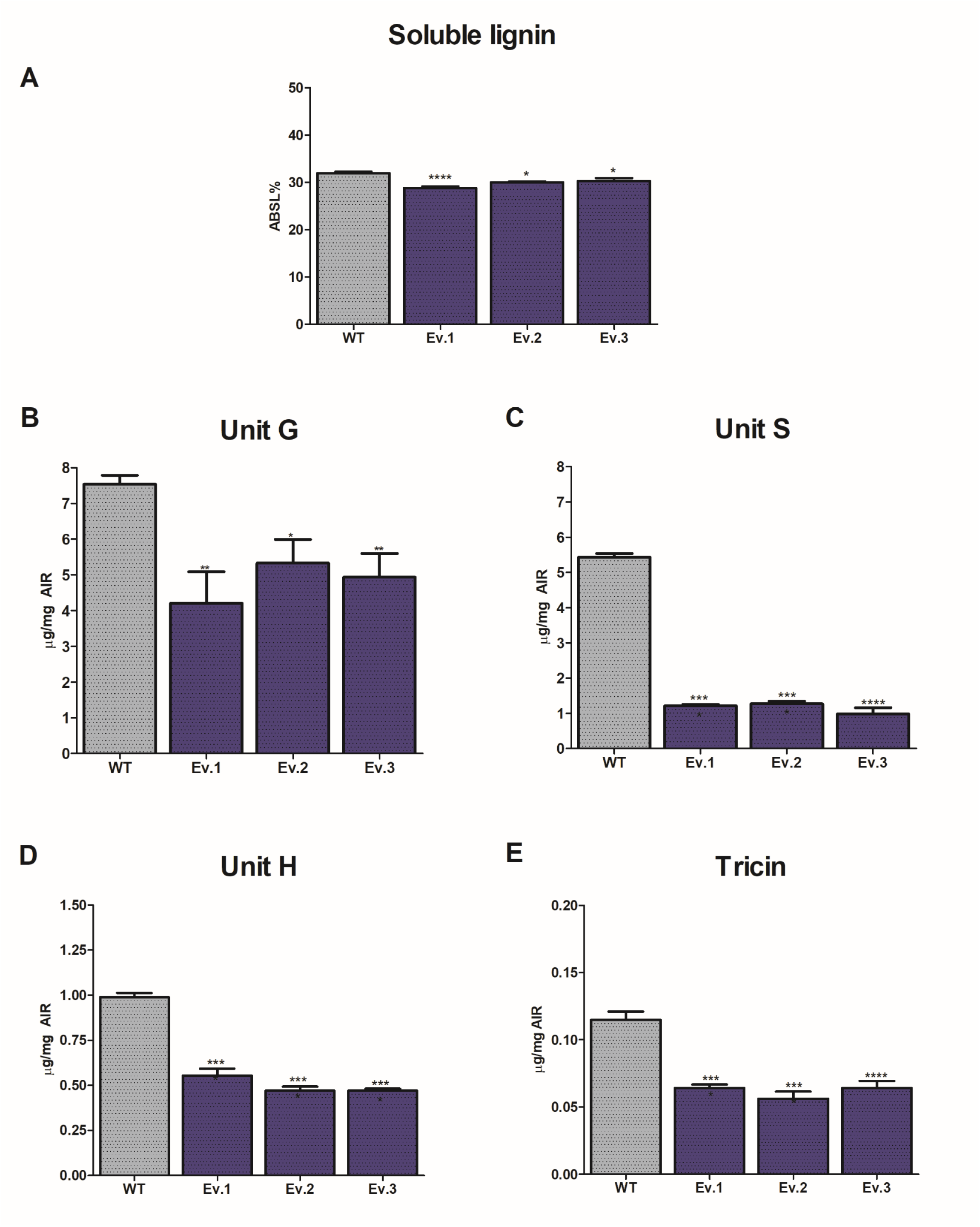
Quantification of lignin content and monomeric composition in wild-type and *COMT1* knockout plants. (A) Soluble lignin content determined by the acetyl bromide method, and levels of (B) guaiacyl (G), (C) syringyl (S), (D) p-hydroxyphenyl (H), and (E) tricin monomers in alcohol-insoluble residue (AIR) extracts. Lignin content and monomeric composition were analyzed in wild-type (WT) plants and in three independent *COMT1* knockout lines (Ev.1, Ev.2, and Ev.3). Values are expressed as μg/mg AIR and represent means ± SEM (n = 4). Statistical differences between WT and knockout lines were assessed using one-way ANOVA followed by Tukey’s post hoc test (*P < 0.05; **P < 0.01; ***P < 0.001; ***P < 0.0001; n.s., not significant).

### 3.4. Total and p-CA-lignin contents are drastically reduced in edited plants

Subsequently, the total content of *p-*CA and FA in *Setaria viridis* was measured by HPLC after alkaline hydrolysis in both mutant and wild-type plants. The method used effectively releases nearly all HCAs that are ester-linked, both in arabinoxylan (AX) and lignin (Möller *et al*., 2022). Our results revealed alterations in the *p-*CA levels in *COMT1* plants. Specifically, as shown in Figure 4A, the total content of *p-*CA decreased by 45% to 51% compared to wild-type plants. Significant changes in FA were also observed in the Ev.2 and Ev.3 lines, with a modest increase of 6% and 4%, respectively. Due to the potential association of ester-linked *p-*CA with both AX and lignin, we employed the methodology described in Lapierre et al. (2018) to extract and quantify HCAs and their conjugates with arabinose (FA-Ara and *p-*CA-Ara) using mild acidolysis. This method allowed us to distinguish between HCAs associated with AX as their arabinose conjugates and those associated with lignin, by subtracting the values of *p*CA-Ara and FA-Ara obtained through mild acidolysis from the values of *p-*CA and FA obtained through alkaline hydrolysis. In this analysis, no significant differences were observed in the levels of *p*-CA and FA esterified to arabinose (*p*-CA-Ara and FA-Ara) in the *COMT1* knockout lines (Ev.1, Ev.2, and Ev.3) compared to the wild-type plants (Figure 4B). However, regarding the *p*-CA bound to lignin, we observed a significant reduction in the *COMT1* knockout plants, with values ranging from 45% to 51%. This decrease was also associated with a 6% and 4% increase in lignin-bound FA in lines #2 and #3, respectively (Figure 4C).

**Figure 4.**
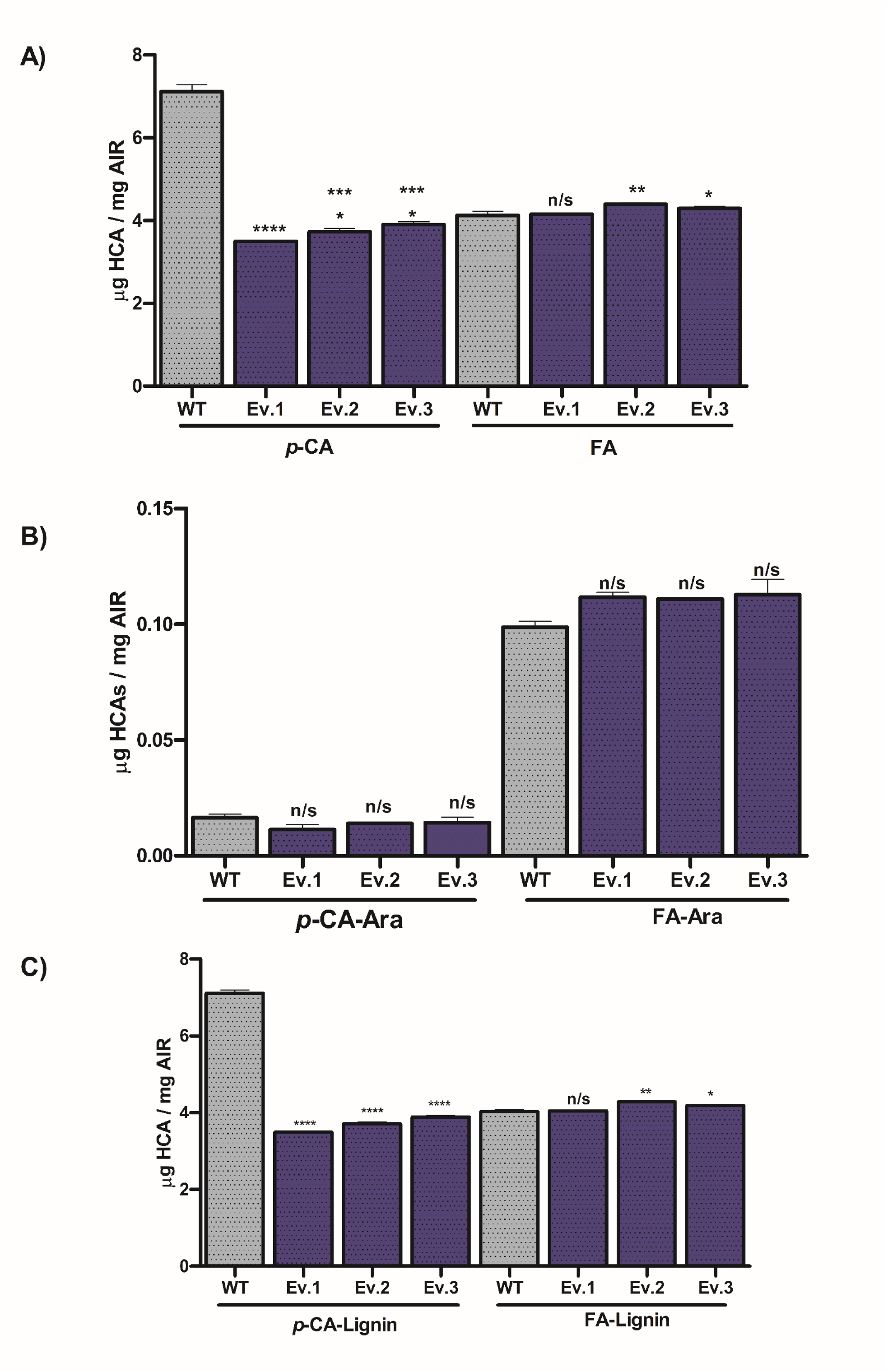
Determination of hydroxycinnamic acid (HCA) content. (A) Total HCA content, (B) HCA content linked to arabinose, and (C) HCA content bound to lignin. The analysis was performed on wild-type (WT) plants and *COMT1* knockout lines (Ev.1, Ev.2, and Ev.3). Values were obtained by alkaline hydrolysis per mg of the AIR (Alcohol-Insoluble Residue) fraction and are expressed as means ± SEM (n = 4). Statistical significance between *COMT1* knockout lines and WT was assessed by one-way ANOVA followed by Tukey’s post hoc test. Significance levels: *P < 0.05; **P < 0.01; ***P < 0.001; ****P < 0.0001; n.s., not significant

### 3.5. Cellulose content and structural monosaccharide composition did not change in mutants compared to wild type plants

The monosaccharide composition of cell wall polymers, determined by high-performance anion-exchange chromatography (HPAEC), showed no significant changes in response to *COMT1* gene knockout (Supplementary Figure 5A). Following this analysis, we evaluated cellulose content, which displayed a slight increase in *COMT1*-knockout plants compared to wild-type controls; however, this difference was not statistically significant (Supplementary Figure 5B). Collectively, these results suggest that the major cell wall modification in the edited plants is associated primarily with alterations in lignin composition.

### 3.6. The enzymatic hydrolysis of biomass revealed that loss of *COMT1* increases the saccharification

To assess the impact of *COMT1* knockout on biomass saccharification, enzymatic hydrolysis assays were performed. Biomass from mutant plants exhibited a significant increase in saccharification efficiency, ranging from 34% to 55% relative to wild-type plants, without pretreatment (Figure 5A). Following pretreatment with 500 mM NaOH, saccharification efficiency increased up to threefold in all lines, likely reaching a sugar release plateau. Although *COMT1* knockout plants showed a modest increase in saccharification compared to wild-type, statistical significance was observed only in line #3 (Figure 5B), with no significant differences detected in lines #1 and #2.

**Figure 5.**
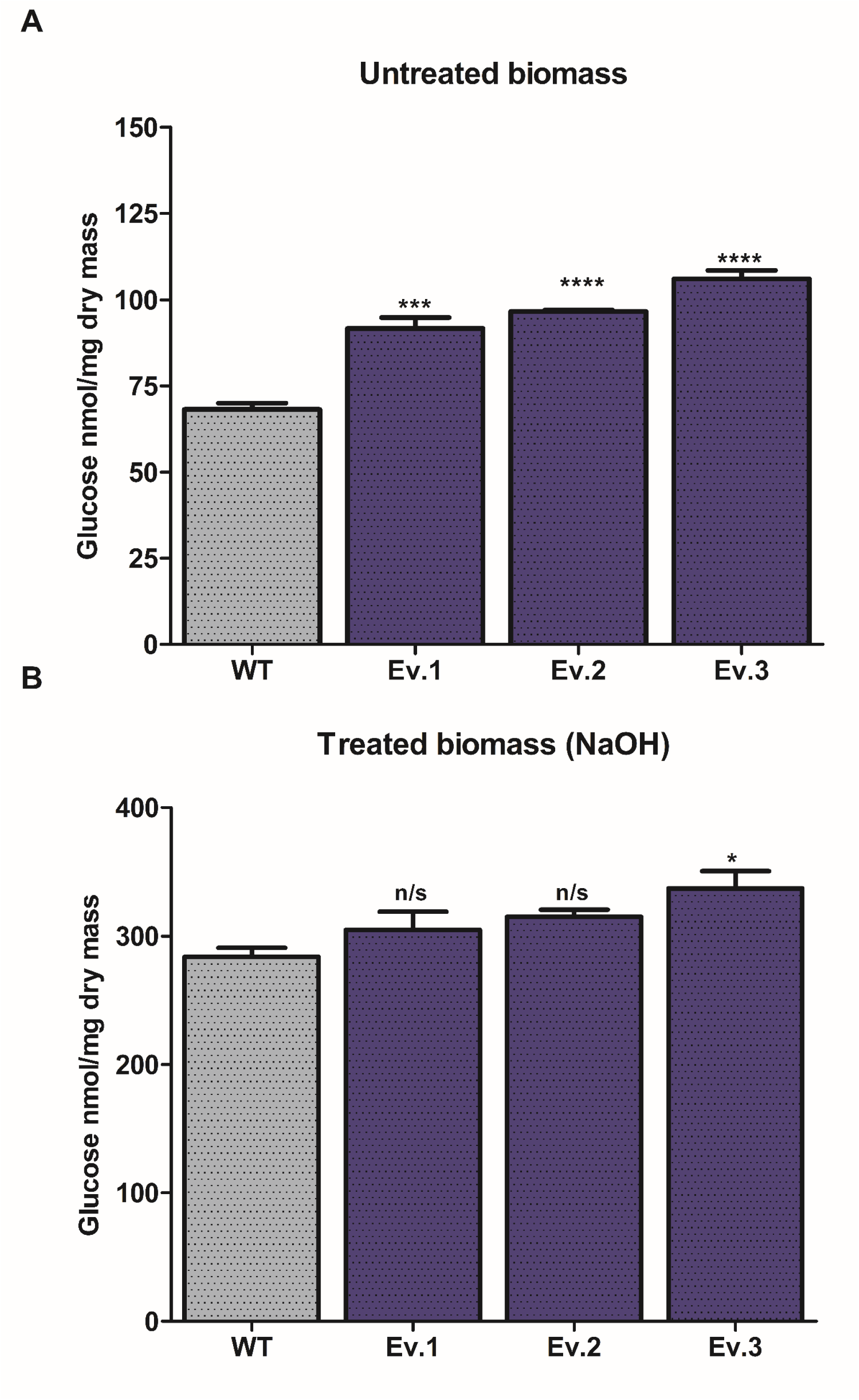
Biomass saccharification analysis. The analysis was conducted on wild-type (WT) plants and *COMT1* knockout lines (Ev.1, Ev.2, and Ev.3) using untreated biomass (A) and biomass treated with 500 mM NaOH (B). Values are expressed as nmol of glucose released per mg of AIR (alcohol-insoluble residue), and data are presented as means ± SEM (n = 4). Statistical significance between *COMT1* knockout lines and WT was assessed by one-way ANOVA followed by Tukey’s post hoc test. Significance levels: *P < 0.05; **P < 0.01; ***P < 0.001; ****P < 0.0001; n.s., not significant.

### 3.7. COMT mutants present altered metabolic flow

Lignin pathway perturbations frequently lead to transcriptional and metabolic reprogramming of the phenylpropanoid pathway, which has been observed for different target genes and plant species (Vanholme *et al*., 2012; Saleme *et al*., 2017; Ferreira *et al*., 2022). Thus, we anticipated that loss of COMT activity results in a redirection of the carbon flux, especially through the phenolic metabolism, given its involvement in different steps of this pathway. To gain insight into the consequences of *COMT1* mutation in the metabolic profile of *S. viridis*, we analyzed acetonitrile-soluble metabolites in aerial parts (leaves + stem) extracts using ultra-high-performance liquid chromatography-mass spectrometry (UHPLC-MS). We were able to detect 956 peaks [mass-to-charge ratio (*m/z*) features] with intensities ≥ 1% of the spectra baseline peak (Supplementary Table 4). Then we filtered these peaks based on blank samples, in which all peaks present in the blank samples were removed from analysis when their intensities were comparable to the plant samples, resulting in 160 peaks (Supplementary Table 5). Next, we focused on peaks present in all replicates of at least one sample type, with at least a twofold difference in intensity in *COMT1* mutants compared with controls, resulting in 22 peaks with higher and lower abundances in mutant plants (Table 1).

**Table 1.**
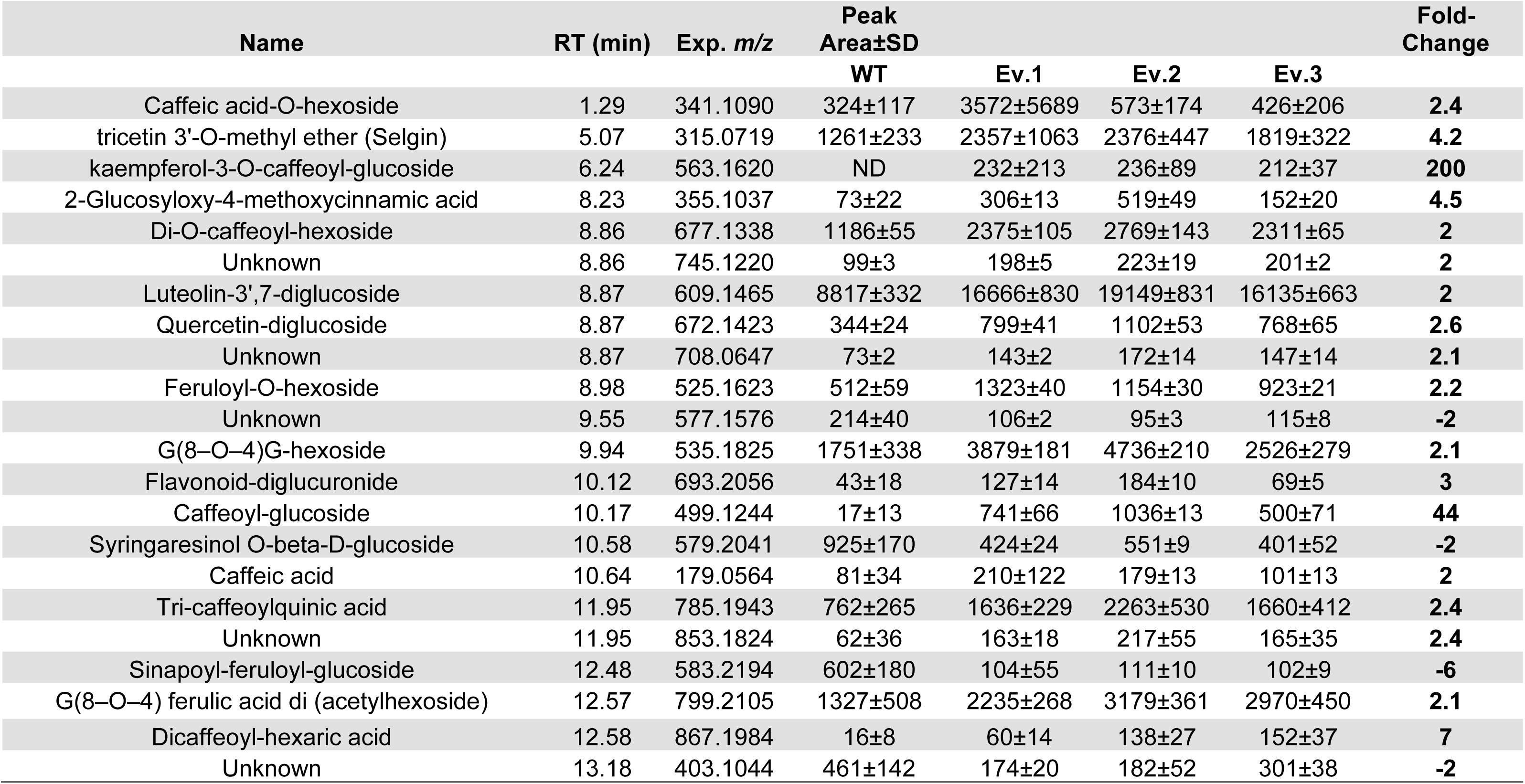
Differentially accumulated compounds in aerial parts of *SvCOMT1* mutants.

From the 22 peaks differentially abundant in *COMT1* mutants, identified 17 peaks (partially) structurally annotated based on tandem mass spectrometry (MS/MS) fragmentation spectra, using different database analysis. Most were phenolic or flavonoid compounds coupled to sugar moieties, such as hexoses and glucuronic acid, including caffeic acid-*O*-hexoside, kaempferol-3-*O*-caffeoyl glucoside, 2-glucosyloxy-4-methoxycinnamic acid, di-*O*-caffeoyl-hexoside, luteolin-3,7-diglucoside, quercetin-diglucoside, feruloyl-*O*-hexoside and caffeoyl-glucoside. Two compounds are putative G-G dimers coupled to a hexoside moiety, G(8–*O*–4)G-hexoside and G(8–*O*–4) ferulic acid di-(acetylhexoside), increased by two-fold in *COMT* mutants. We also observed one compound involved in phenylpropanoid biosynthesis directly linked to lignin, caffeic acid, increased by 2-fold in *COMT1* mutants. From these 22 peaks, only 4 were decreased in mutant plants, syringaresinol *O*-beta-D-glucoside (decreased by 2-fold), sinapoyl-feruloyl-glucoside (decreased by 6-fold), and 2 unidentified compounds, with *m/z* 577.1576 and 403.1044 and retention time 9.55 and 13.18, respectively. These results suggest that disruption of *COMT1* leads to rewiring of the phenolic metabolism in *S. viridis*.

### 3.8. *COMT1* knockout plants did not present altered sensitivity to moderate or severe salinity stress

Genetic modifications in the plant cell wall can result in pleiotropic effects, including altered responses to abiotic stresses such as salinity. To determine whether *COMT1* knockout influences plant tolerance to salt stress, a major environmental factor limiting growth and agricultural productivity, we evaluated *COMT1* knockout lines under different NaCl concentrations. Wild-type (WT) plants and knockout lines (Ev.1, Ev.2, and Ev.3) were exposed to 0, 100, and 250 mM NaCl for 5 and 21 days. These NaCL concentrations have been previously shown to cause stress responses in S. viridis (Duarte *et al*., 2019). Representative phenotypes after 5 days across all treatments are shown in Figures 6A and 6B, while phenotypes after 21 days are presented in Supplementary Figure 6. At 250 mM NaCl, plants of all genotypes failed to survive beyond 9 days (Figure 6C). Conversely, plants submitted to 100 mM NaCl, despite a slight decrease in biomass compared to control (Figure 6B), did not present differences in relative water content (Figure 6A), suggesting a moderate tolerance to salt stress. As shown in Figure 6, this tolerance was not affected by mutations in *COMT1*, as no significant differences in relative water content (RWC) or dry mass were detected between WT and knockout lines after 5 days of 100 mM NaCl (Figures 6D and 6E) or 21 days under 100 mM NaCl (Supplementary Figure 6). These results indicate that *COMT1* loss does not impair growth or salt tolerance, as knockout plants maintained RWC levels comparable to WT under saline conditions.

**Figure 6.**
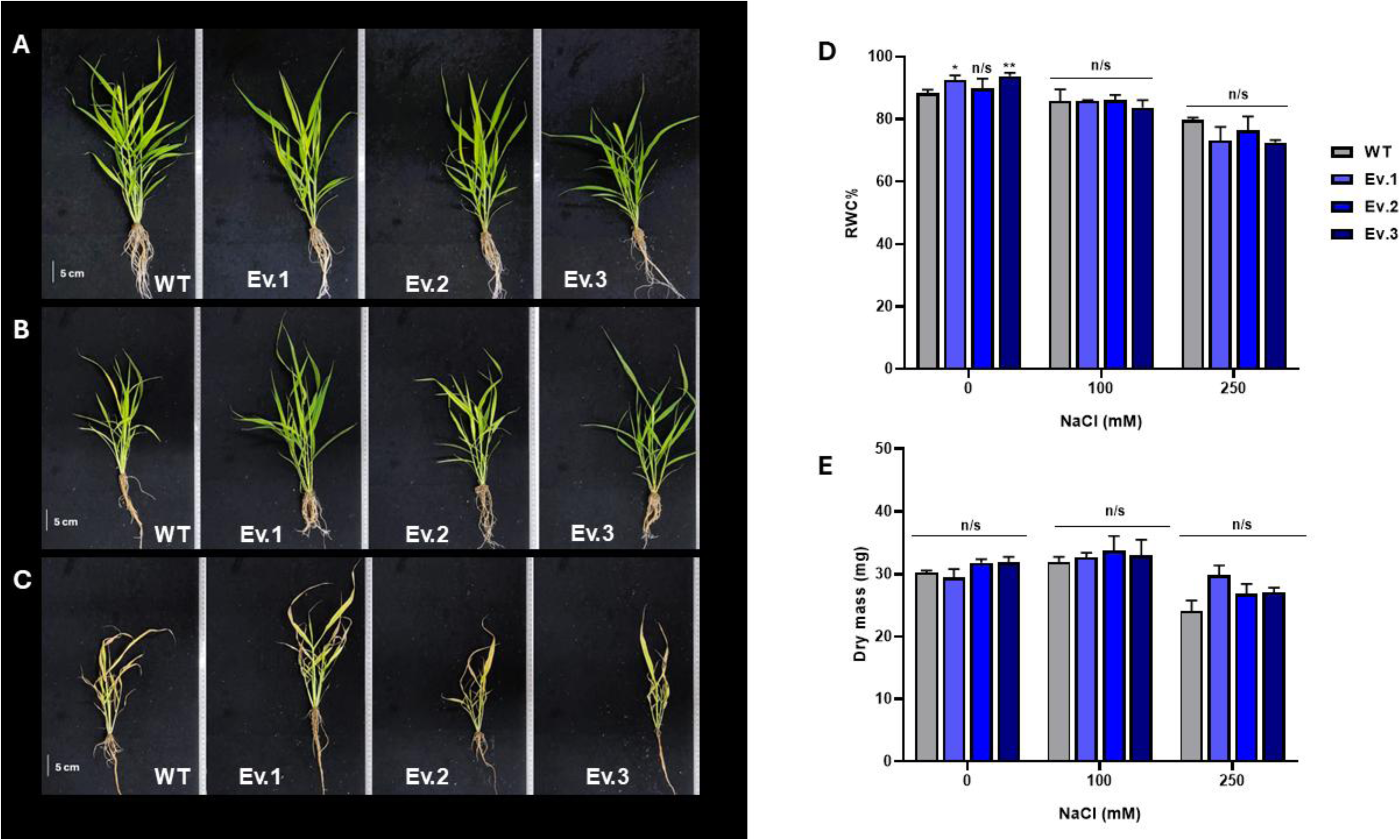
Responses of Relative Water Content and Dry Mass in Wild-Type and *COMT1* Knockout Plants to 5-Day NaCl Treatments. (A–B) Representative phenotypes after 5 days of exposure to 0, 100, and 250 mM NaCl. (C) Survival of plants exposed to 250 mM NaCl, showing that all genotypes failed to survive beyond 9 days. (D–E) Relative water content (RWC) and dry mass after 5 days under the indicated NaCl concentrations. Values are presented as means ± SEM (n = 4). Statistical significance relative to WT was determined by one-way ANOVA followed by Tukey’s post hoc test (p ≤ 0.05); n.s., not significant.

## 4. Discussion

The conversion efficiency of lignocellulosic biomass into biofuels is inversely correlated with the lignin content in feedstock plants, as demonstrated by Chen and Dixon (2007) and Studer et al. (2011), among many other studies. This correlation arises from the recalcitrance imposed by lignin in the cell wall, which limits enzyme access to structural polysaccharides. Consequently, strategies aimed at reducing lignin content or modifying its structure have been extensively explored to improve biofuel yields from lignocellulosic biomass (Dixon *et al*., 2001; Fu *et al*., 2011; Baxter *et al*., 2014). Different genetic engineering approaches have been employed to decrease lignin content and alter its composition in different species. However, disrupting lignin biosynthesis can lead to undesirable pleiotropic effects, such as reduced mechanical strength and decreased stem rigidity. For instance, in *Arabidopsis thaliana*, mutants with a 50% reduction in lignin content exhibit weaker stems (Jones *et al*., 2001). Likewise, in tobacco, downregulation of an *O-methyltransferase* reduces lignin levels but also negatively impacts plant growth (Pichon *et al*., 2006).

It is currently known that COMT enzymes play a crucial role in lignin biosynthesis in angiosperms by catalyzing the methylation of the hydroxyl group at the 5-position (5-OH) of intermediates in the phenylpropanoid pathway (Bonawitz and Chapple, 2010, Preprint; Vanholme *et al*., 2010). This process is essential for the formation of sinapyl alcohol, the main precursor of the S lignin units (Figure 1). Studies indicate that the primary *in vivo* substrates for COMTs are 5-hydroxyconiferaldehyde and 5-hydroxyconiferyl alcohol, intermediates required to produce sinapyl alcohol (Zubieta *et al*., 2002; Louie *et al*., 2010; Green *et al*., 2014). In addition to the methylation of S unit precursors, it is currently accepted that COMT is a bifunctional enzyme in lignin biosynthesis, at least in grasses. In this context, COMT is also responsible for methylation steps that lead to the production of tricin, a member of the flavone subclass of flavonoid compounds, which is incorporated into the lignin polymer via 4-O-β-coupling and is considered the first lignin non-canonical monomer outside the monolignol biosynthetic pathway (Del Río *et al*., 2012; Lan *et al*., 2016*a*,*b*).

In *Setaria viridis*, a study conducted by Ferreira et al. (2019) six *COMT* genes were identified in its genome, all belonging to class II of the *O-methyltransferase* superfamily. These *COMTs* form a distinct phylogenetic group within the monocots, with *COMT1* and *COMT6* showing significant similarity to *COMT* genes involved in lignification in grasses such as *Brachypodium distachyon*, maize, rice, and sorghum. The strong expression of *COMT1* in the transition and maturation zones of the elongated internode in *S. viridis* suggests a central role in lignification, making it an ideal target for genetic manipulation to reduce biomass recalcitrance and improve saccharification efficiency. Therefore, we have chosen *SvCOMT1* as a target for the generation of CRISPR/Cas9 mutants in *S. viridis* to disrupt this gene.

### The disruption of *COMT1* gene in *Setaria viridis* did not show any visible phenotype

In some C4 grasses, such as *Zea mays* and *Sorghum bicolor*, mutations in genes involved in lignin metabolism, including *COMT*, result in the brown midrib phenotype (*bm* or *bmr*), which is characterized by brown-colored central veins and, often, other parts of the plants. This phenotype is associated with reduced lignin content and increased biomass digestibility (Vignols *et al*., 1995; Bout and Vermerris, 2003), and it is widely used to identify mutants affected in lignin deposition (Sattler *et al*., 2010, Preprint). Whereas the majority of the isolated mutants with this phenotype have been characterized as affected in lignin biosynthesis, loss-of-function of genes encoding enzymes that synthesize the methyl donor *S*-adenosyl-L-methionine, used by lignin *O*-methyltransferases, was also shown to result in brown midribs in maize (Tang *et al*., 2014; Li *et al*., 2015). Additionally, the sorghum *bmr30* mutant has been characterized as disrupted in the tricin biosynthetic gene encoding chalcone isomerase (Tetreault *et al*., 2021). Until recently, brown midribs have been described only in a few lignin mutants of panicoid C4 grasses such as maize and sorghum, but disruption of some lignin biosynthetic genes in the C3 grass *Brachypodium distachyon*, including *COMT*, also resulted in the accumulation of brown pigment in leaf midribs and other lignified tissues (Bouvier D’Yvoire *et al*., 2013; Trabucco *et al*., 2013*b*). Conversely, this phenotype has not been found in rice or barley, despite their extensive mutant stocks. Regarding *COMT*, their corresponding mutants in maize and sorghum, as well as their silenced lines in Brachypodium and switchgrass, showed the brown midrib phenotype (‘Bout 2003; Piquemal *et al*., 2002*b*; Trabucco *et al*., 2013*b*; Wu *et al*., 2019). In addition to the *bmr* phenotype, different studies demonstrated that null *comt* mutant plants presented an abnormal growth (Yu *et al*., 2025), while silenced *COMT* gene plants have reduced lignin content without growth penalties (Vermerris *et al*., 2007; Saballos *et al*., 2008; Dien *et al*., 2009; Fu *et al*., 2011; Jung *et al*., 2012, 2013; Lam *et al*., 2019*a*). Here, the *COMT1* edited *S. viridis* plants did not display any visible phenotype, neither the brown midrib trait, nor observable changes in plant growth (Figure 2). In the context of generating optimized feedstock for bioeconomy, the lack of detrimental effects in plant growth and development is a desirable trait. A similar phenotype was found when more than 100 *COMT* copies/alleles were gene-edited in sugarcane using TALEN technology (Kannan *et al*., 2018), demonstrating that targeting *COMT* genes using gene editing might be an effective strategy for biomass optimization without yield penalty.

### *COMT1* mutants display a drastic reduction of S monomers and a decrease of G, H and tricin moieties in the lignin portion of *S. viridis*

Our results demonstrated a drastic reduction (from 40 to 80% reduction) in S units and tricin in the AIR of edited *COMT1* plants, while G and H monomers demonstrated less pronounced effects (Figure 4). It is expected that disruption of COMT activity leads to reduction of S units, as the enzyme is mostly involved in the pathway steps that generates S monomers in grasses. The novelty here is that tricin, G and H monomers were also significantly decreased, despite the total soluble lignin content only slightly reduced (Figure 4A). These results may indicate that other steps within the phenylpropanoid/flavonoid pathways might be compensating for lignin biosynthesis through the incorporation of non-canonical monomers, to the extent that normal plant development remains unaffected. The fact that total soluble lignin was only slightly decreased may reinforce the speculation that more soluble, non-canonical monomers, could be incorporated into the lignin.

Although the mechanisms underlying lignin homeostasis remain unclear, previous research suggests that plants can preserve lignin deposition by incorporating non-canonical aromatic monomers when canonical monolignols are limited. Reports of flavonoid-derived units, hydroxycinnamate amides, and other atypical phenolics support the view that lignin is more flexible than previously assumed, providing compensatory routes that may stabilize lignin levels despite major pathway disruptions (Ralph et al., 2001b,a; Bonawitz and Chapple, 2010; Vanholme et al., 2010). Also, it is worth mentioning that reduction in S, G and H monolignols may reflect both a global down-regulation of phenylpropanoid flux through feedback inhibition (see Table 1) or/and an altered lignin architecture with increased condensed linkages (C-C linkages instead of β-O-4), which are poorly released by degradative methods.

Although there are studies that partially characterize lignin biosynthesis in plants such as *Arabidopsis*, *Oryza sativa*, and *Zea mays*, there are still significant gaps in our understanding of the dynamics of this process in grasses. For example, recent research indicates that many stages of monolignol biosynthesis in grasses do not occur in the same way as in eudicots. In *Arabidopsis*, loss-of-function mutations in genes such as *F5H* and *COM*T lead to a significant reduction in sinapyl lignin units, resulting in a polymer composed solely of G units (Meyer *et al*., 1998; Goujon *et al*., 2003). However, in *Oryza sativa*, the loss of function of the *F5H* gene does not result in a complete absence of S units, while in *COMT*-deficient mutants of maize and sorghum, S units are significantly reduced but not completely absent. This suggests that grasses may have alternative pathways to produce these units (Piquemal *et al*., 2002*a*; Fornalé *et al*., 2017; Takeda *et al*., 2019). Our results on lignin monomers align with those previously reported in the literature for *Brachypodium distachyon*, *Zea mays*, and *Hordeum vulgare* with *COMT* silencing (Piquemal *et al*., 2002*a*; Barrière *et al*., 2004; Daly *et al*., 2019). However, these studies reported that the reduction in S lignin content does not affect G monomers, which contrasts with the findings of the present work. As mentioned above, in *S. viridis* the *COMT1* knockout resulted in significant reductions in G lignin units (ranging from 13.3% to 28.8%), contrary to what was reported in those studies. In alfalfa plants, the negative regulation of *COMT* also caused a decrease in G units and an almost complete loss of S units (Guo *et al*., 2001). While the formation of G lignin units predominantly depends on CCoAOMT activity, *COMT* may play a secondary role by contributing to the methylation of intermediates feeding into this pathway, such as coniferaldehyde. Thus, the absence of *COMT* could reduce the efficiency of the G unit formation pathway, which explains the decrease in G lignin units observed in our study. Moreover, the reduction in H units observed in *S. viridis* (ranging from 44.1% to 52.5%) can be attributed to the overall modulation of the lignin biosynthetic pathway, where disruption of methylation affects both the accumulation and utilization of precursor monomers. These combined effects confirm that the knockout of the *COMT1* gene not only drastically reduces S units but also impacts the synthesis of other lignin monomers.

### *COMT1* mutants demonstrate substantial differences in *p*CA pools

In grasses, lignin and xylan polymers can be substituted with hydroxycinnamic acids (HCAs), such as ferulic acid (FA) and *p-*coumaric acid (*p-*CA). These HCA groups can be esterified or etherified in lignin, playing a crucial role in modulating the structure and properties of the cell wall (Hatfield *et al*., 2017). In particular, the feruloylation of xylan facilitates the formation of covalent cross-links between xylan chains or between xylan and lignin through the formation of a diferulate (diFA) bridges. This cross-linking process contributes to the rigidity and mechanical strength of the cell wall, as well as affecting biomass digestibility (Hatfield *et al*., 2017; Terrett and Dupree, 2019, Preprint; Feijao *et al*., 2022).

Recent studies suggest that, in grasses, the phenylpropanoid pathway is predominantly directed toward the synthesis of hydroxycinnamic acids (HCAs) (Barros et al. 2022; Karlen et al. 2018; Ralph 2010). This pattern can be attributed to the higher abundance of *p*-coumaric acid (*p*-CA) and ferulic acid (FA) in the cell walls of grasses compared to dicotyledonous species. In this context, *p*-CA plays a crucial role as a precursor in the biosynthesis of various phenylpropanoids and flavonoids (Vogt, 2010) and is commonly conjugated to syringyl (S) and guaiacyl (G) lignin monomers (Peracchi *et al*., 2024, Preprint). Although the exact function of *p*-CA–lignin conjugates is not fully understood (Hatfield et al. 2017), two main hypotheses have been proposed: (1) *p*-CA may act as a radical transfer agent during the polymerization of S units (Ralph *et al*., 2004), or (2) *p*-CA-S monomers could terminate polymerization, resulting in a more linear lignin polymer and consequently altering its structural properties (Kishimoto *et al*., 2005; Hatfield *et al*., 2017). In addition to its conjugation with S lignin monomers, *p*-coumaric acid (*p*-CA) is also acylated to arabinoxylans in the cell walls of grasses. This process is mediated by BAHD acyltransferases, as demonstrated in species such as *Oryza sativa* (Bartley *et al*., 2013) and *Setaria viridis* (de Souza *et al*., 2018; Mota *et al*., 2021).

In the present study, using alkaline hydrolysis and mild acidolysis, we were able to quantify total *p*CA and FA, in addition to *p*CA-Ara and FA-Ara, respectively. Further calculations allowed us to estimate the amount of *p*CA and FA bound to lignin (Figure 4). We observed that there was a large reduction of total *p*CA and *p*CA bound to lignin, while *p*CA-Ara did not show any significant changes in *COMT1* mutants. These results indicate a substantial modulation in the *p*-CA pools, demonstrating that the knockout of *COMT1* directly affects the incorporation of *p*-CA into cell walls, thereby influencing lignin formation and structure. The reduction in total and lignin-bound *p*-CA content observed in our study is consistent with previous reports investigating negative regulation of *COMT* in maize (Pichon *et al*., 2006) and *Brachypodium distachyon* (Ho-Yue-Kuang *et al*., 2016), in which decreased levels of *p*-CA and S-type lignin monomers were also observed. Moreover, our results reveal a slight increase in total FA content (ranging from 4% to 6%) in *COMT1* knockout plants. This increase may be associated with compensatory adjustments in the phenylpropanoid biosynthesis caused by disruption of *COMT1*. Evidence of this mechanism has already been reported in genetic modification studies involving the addition of HCAs in *Oryza sativa* and *Setaria viridis* (de Souza *et al*., 2018; Möller *et al*., 2022). Therefore, we propose that in *S. viridis* ferulic acid biosynthesis is not directly impacted by the knockout of *COMT1*, as its production may proceed via an alternative pathway that bypasses caffeic acid methylation, as suggested by previous studies (Barrière *et al*., 2004; Chen *et al*., 2006).

### *COMT1* disruption increased biomass saccharification without altering cell wall polysaccharides

*COMT1* knockout revealed no significant changes in the contents of structural polysaccharides (Supplementary Figure 5A). This suggests that, while *COMT* plays a key role in lignin biosynthesis, it does not directly affect cell wall polysaccharides. This result is supported by studies in Arabidopsis and barley, in which loss-of-function of *COMT* did not lead to changes in cellulose or structural carbohydrate content (Lee *et al*., 2021). Conversely, the amounts of arabinose and xylose released from hemicellulosic polymers were significantly increased in both *COMT* downregulated and knockout rice plants (Lam *et al*., 2019*b*), demonstrating different results depending on the plant species. As expected for plants with diminished lignin contents, the *COMT1* knockout plants showed a significant increase in biomass saccharification efficiency, ranging from 34% to 55% compared to WT plants, without the necessity for pretreatments (Figure 5A). When subjected to pretreatment with 500 mM NaOH, saccharification increased up to threefold, regardless of the genotype, potentially reaching a plateau in sugar release. However, the *COMT1* knockout plants showed only a slight increase in saccharification efficiency compared to the WT plants upon alkaline pretreatment, with no statistically significant differences (Figure 5B). The absence of significant differences after pretreatment may be explained by the effectiveness of the pretreatment in breaking down the structural bonds of the cell wall, making the biomass more accessible to enzymes and masking potential differences between the groups. Our results are in line with previous studies that reported a negative correlation between lignin content and the efficiency of hemicellulose enzymatic hydrolysis and fermentation in species such as switchgrass, sugarcane, sorghum, and alfalfa (Chen and Dixon, 2007; Fu *et al*., 2011; Xu *et al*., 2011; Jung *et al*., 2012).

### Metabolome profiling of *COMT1* mutants suggests a redirection of carbon flux towards phenylpropanoid and flavonoid pathways

Our metabolome analysis demonstrated that, as expected, *COMT1* disruption led to accumulation of caffeic acid, a known (but not preferential) substrate of COMT enzymes (Table 1). However, most tentatively annotated compounds accumulating in *COMT1* mutants were conjugated forms of either flavonoids or phenylpropanoids, such as kaempferol-3-*O*-caffeoyl-glucoside (200-fold increase), 2-glucosyloxy-4-methoxycinnamic acid (4.5-fold increase), caffeoyl-glucoside (44-fold increase) and quercetin-diglucoside (2.6-fold increase). It is not surprising that most of the accumulated metabolites found in edited plants were found as their sugar-conjugated forms, given that several phenylpropanoid derivatives are either cytotoxic and/or bioactive and glycosylation is known to play key roles in modulating phenylpropanoid properties (including solubility, toxicity and bioactivity) and compartmentalization (Rates and Cesarino, 2023, Preprint).

Our results are consistent with some recent studies showing that perturbations of lignin metabolism in grasses lead to the accumulation of glycosylated phenylpropanoids (Barros *et al*., 2022; Ferreira *et al*., 2022; Oliveira *et al*., 2025; Priya *et al*., 2025, Preprint). Importantly, we observed that selgin, the direct precursor of tricin biosynthesis and a known substrate for grass COMTs, is highly accumulated in mutant plants (4.5-fold increase). In addition, luteolin, which is also an intermediate of the tricin pathway and a COMT substrate, also accumulated, but in its conjugated form as luteolin-3’,7-diglucoside (2-fold increase). The fact that we did not find in our metabolome many intermediates of the phenylpropanoid pathway suggests that *S. viridis* may have other OMTs involved in lignin biosynthesis through this pathway, or the disruption of *COMT1* is redirecting the carbon flux to flavonoid synthesis. Thus, our results further corroborate that *Setaria viridis* COMT1 play an important role in the synthesis of tricin. The accumulated flavonoids could also be responsible for the resilience of the mutants under salt stress observed in our work.

### Salinity stress tolerance is not altered in *COMT1* mutants

Lignin plays a crucial role in plant defense against pathogens and in adaptation to stresses, such as salinity. Given the central role of the *COMT* gene in lignin biosynthesis, we investigated how its knockout affects plant responses to abiotic stresses, particularly salt stress. Soil salinization is a major form of land degradation, characterized by the excessive accumulation of water-soluble salts, especially sodium (Na⁺) and chloride (Cl⁻) ions (Tian *et al*., 2021). Several studies have demonstrated that the *COMT* gene regulates plant stress responses, including salinity (Sun *et al*., 2020; Wang *et al*., 2024), cold (Chang *et al*., 2021), and drought (Yao *et al*., 2022).

Interestingly, although previous studies have demonstrated that genetic modifications in the *COMT* gene can influence plant responses to salinity, our results show that under moderate salinity levels (100 mM), not only the WT but also genetically modified *Setaria viridis* plants are able to maintain relative water content (RWC) in their tissues (Figure 6), suggesting a moderate tolerance of *S. viridis* to salt stress. Under more severe salinity conditions (250 mM), a reduction of 10.8% in RWC was observed only in line Ev.3, whereas the other edited lines did not show significant changes. Moreover, plant survival was not affected compared to WT, as both WT and *COMT*-knockout plants were able to survive for up to nine days under high salt concentrations. Furthermore, dry biomass measurements (Figure 6) revealed no statistically significant differences between WT and *COMT*-knockout plants at any of the tested NaCl concentrations, both after 5 and 21 days of treatment. These findings suggest that *COMT* gene editing does not significantly affect biomass accumulation under the salinity conditions evaluated.

In conclusion, this study highlights the potential of CRISPR/Cas9-mediated knockout of the *COMT1* gene as an effective strategy to alter lignin and improve biomass digestibility in grasses. This has direct implications for enhancing saccharification efficiency and, consequently, biofuel production, without increasing the plants’ susceptibility to salt stress. Although the typical *bmr* phenotype was not observed in the mutant lines, the changes in lignin composition and the significant reduction in S, G, H, and tricin monomers reinforce the role of *COMT1* in modulating lignin structural diversity in grasses. Moreover, the alterations in *p*-coumaric acid (*p*-CA) and ferulic acid (FA) content, combined with metabolomics, suggest the activation of a compensatory mechanism that maintains cell wall structural integrity under both normal and saline conditions. In this context, the application of techniques such as nuclear magnetic resonance (NMR) spectroscopy would be valuable for gaining deeper insights into how *COMT1* knockout-driven modifications affect lignin structure, especially regarding incorporation of non-canonical monomers. Such information could significantly contribute to the rational engineering of plant cell walls without compromising growth. These findings support the notion that genetic manipulation of lignin biosynthetic genes represents a promising approach for optimizing lignocellulosic biomass conversion, with applications in bioenergy and animal nutrition, offering a meaningful contribution toward the development of more efficient and sustainable renewable energy technologies.

## Author Contributions

**Fernanda de Oliveira Menezes:** methodology; investigation; formal analysis; data curation; writing original draft; writing review & editing. **Karoline Estefani Duarte:** supervision; writing review & editing. **Gabriel Garon Carvalho:** formal analysis. **Leonardo Dario Gomez:** methodology; supervision; writing review & editing. **Igor Cesarino:** formal analysis; writing review & editing. **Wagner Rodrigo de Souza:** conceptualization; supervision; project administration; funding acquisition; writing review & editing.

## Conflict of Interest

No conflict of interest declared.

## Funding Statement

This research was supported by the Fundação de Amparo à Pesquisa do Estado de São Paulo (Fapesp Proc. 2022/04419-5, 2019/26761-4 and 2019/04878-7).

## Notes

### Competing Interest Statement

The authors have declared no competing interest.

### Summary of Updates

The revised version includes Supplementary Figures S1 to S6 and and Suppl Tables S1 to S5

