## Supplementary Figures for "Gene Editing of Caffeic-O-methyltransferase (COMT1) in the model grass Setaria viridis Improves Biomass Saccharification Without Compromising Plant Growth or Abiotic Stress Tolerance"

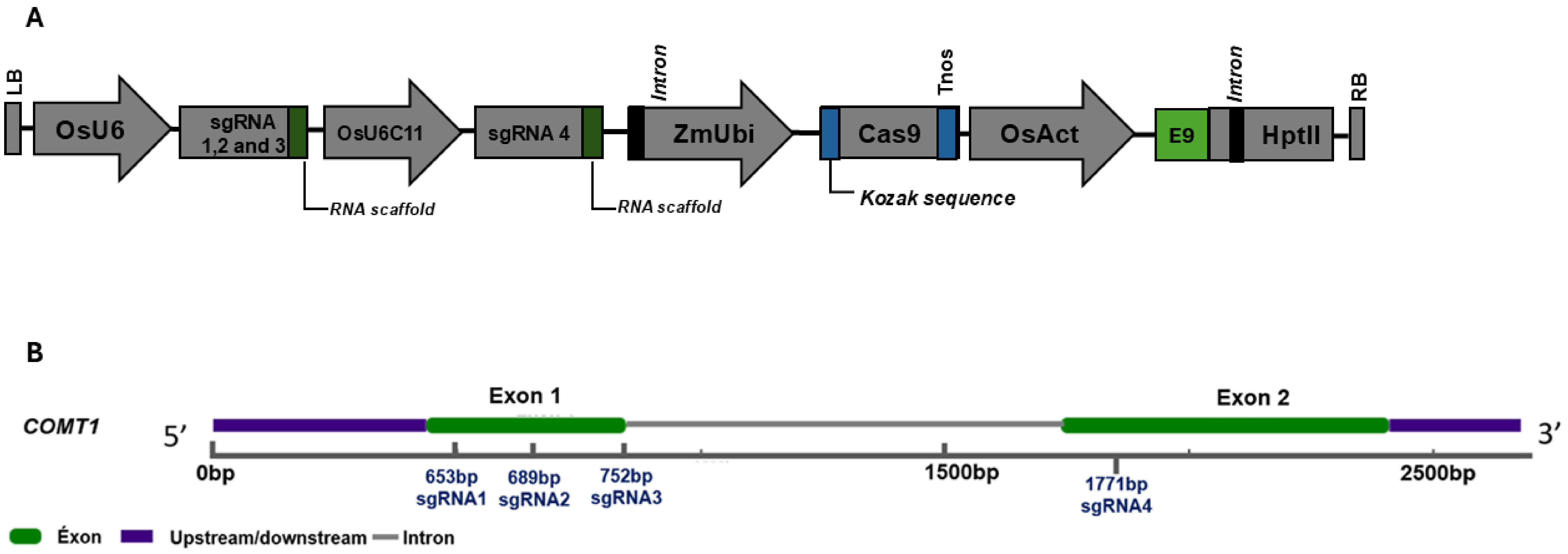


**Supplementary Figure 1. Schematic overview of the optimized CRISPR/Cas9 vector E856 and sgRNA target sites in the *COMT1* gene.**

(A) Map of the CRISPR/Cas9 binary vector constructed for targeted knockout of the *COMT1* gene. Key components include: LB (left border) and RB (right border) sequences; (I) the hygromycin phosphotransferase II (HptII) selection marker gene, containing an intron and driven by the constitutive *Oryza sativa* Actin 1 promoter (OsAct-1) with the 3′ rbcsE9 terminator; (II) the Streptococcus pyogenes wild-type Cas9 (SpCas9WT), codon-optimized for monocots and fused with dual SV40 nuclear localization signals (NLS) at both 5′ and 3′ ends, expressed under the constitutive *Zea mays* Ubiquitin promoter (ZmUbi) containing a monocot-optimized Kozak sequence (CCGAA) immediately upstream of the ATG start codon; (III) a guide RNA (gRNA 1–4) expression cassette, each driven by the constitutive *Oryza sativa* U6 (OsU6) promoters and transcribed by RNA polymerase III, with a supplementary 3′ guanine (G) nucleotide fused to the synthetic RNA scaffold to enhance Cas9 binding.
(B) Schematic representation of the four single guide RNA (sgRNA) target sites distributed along the COMT1 coding sequence.

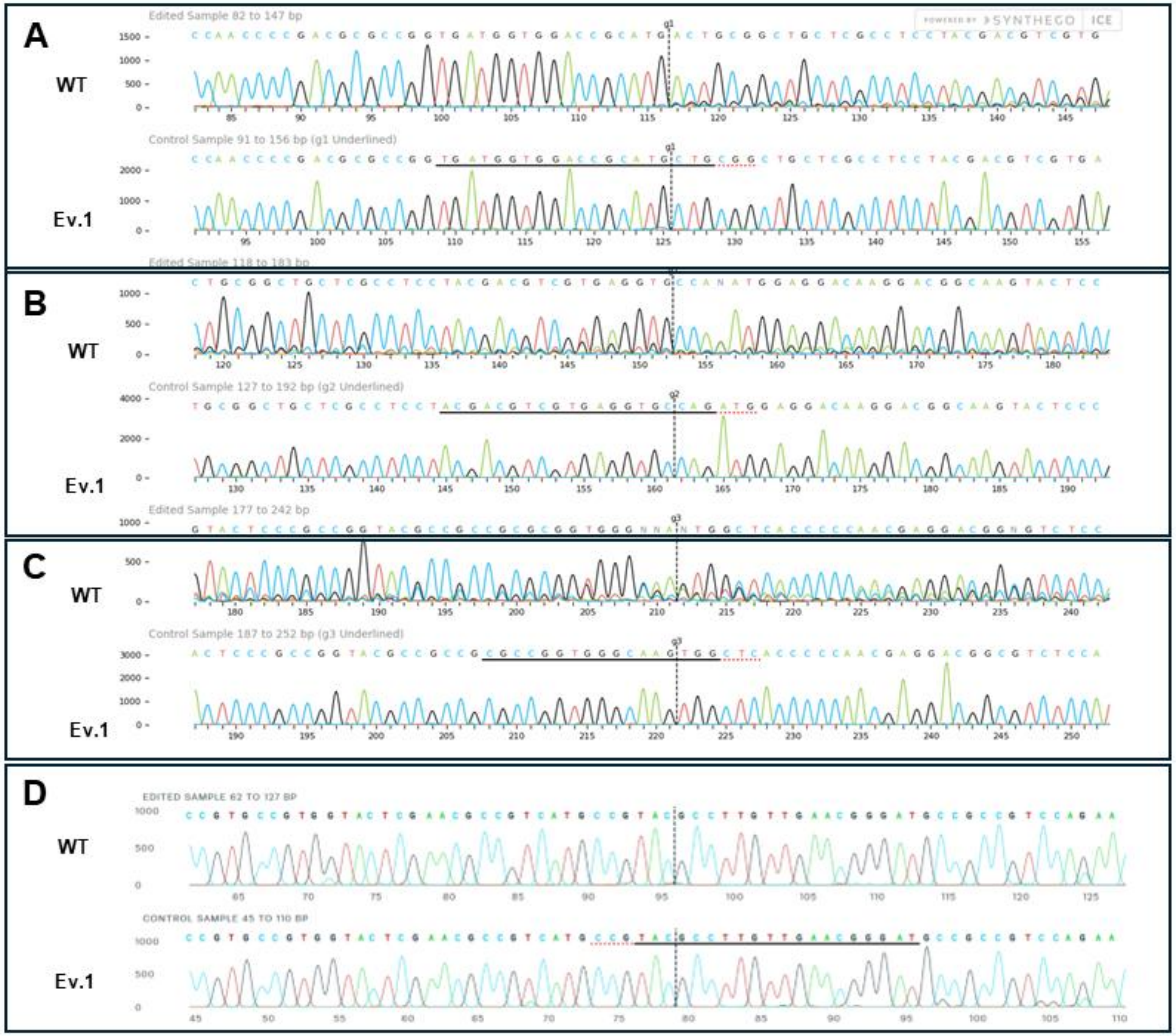


**Supplementary Fig. 2. Sanger Sequencing Chromatogram Analysis of *COMT1* Knockout (Ev.1) Using SYNTHEGO Software**

Fragments A–D correspond to the amplified regions containing the target sites of sgRNAs 1, 2, 3, and 4, respectively. Sanger sequencing results show both edited (Ev.1) and wild-type (WT) sequences, with a particular focus on the region surrounding the guide sequence. Mixed bases are frequently observed, indicating the presence of edited alleles with insertions and/or deletions compared to the wild-type allele. The guide sequence is highlighted with a black underline, while the PAM site is marked with a red underline. The expected Cas9 cleavage site is indicated by a vertical black dashed line, where error-prone repair processes often result in the mixed base signals observed after cleavage.

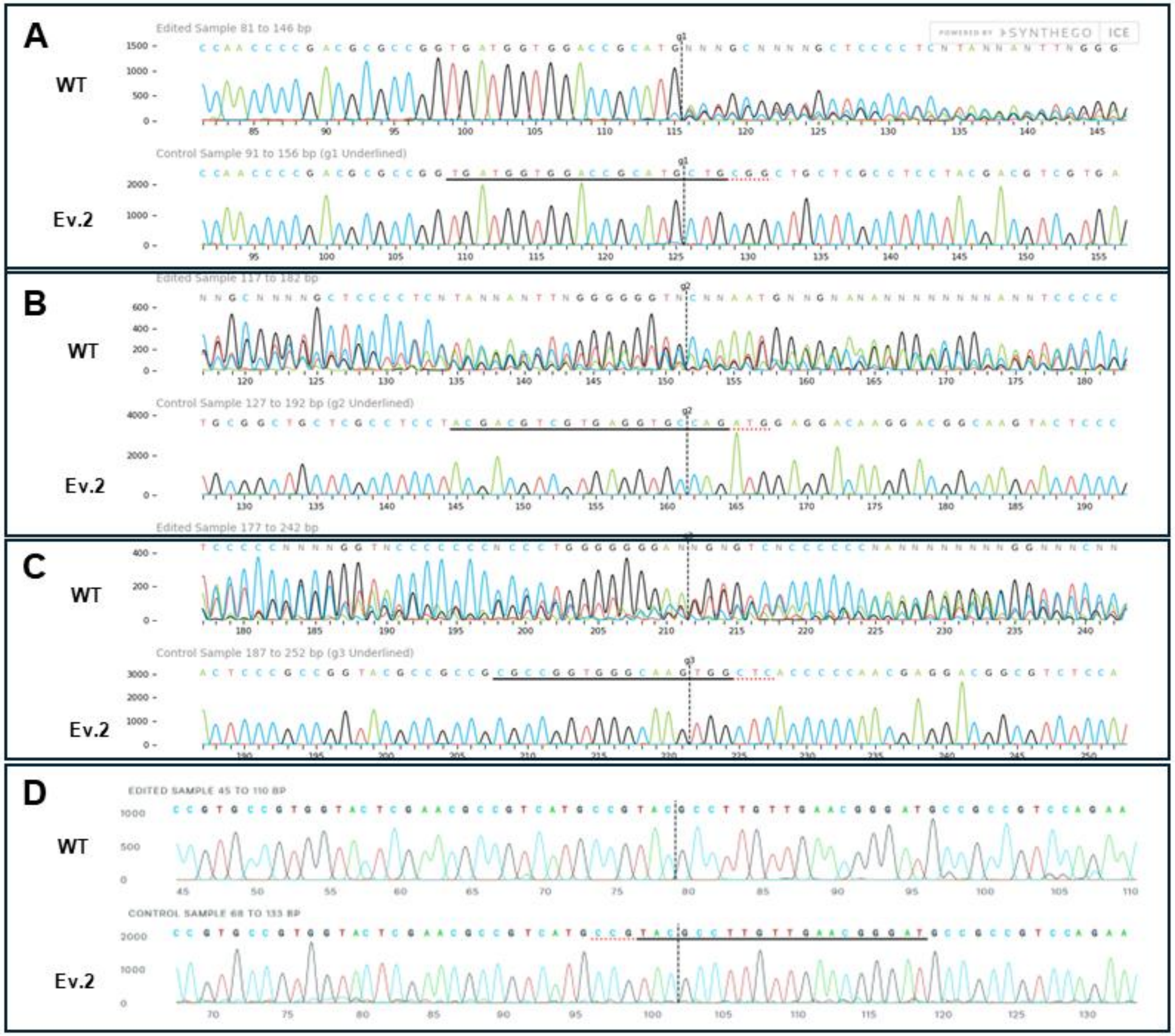


**Supplementary Fig. 3. Sanger Sequencing Chromatogram Analysis of *COMT1* Knockout (Ev.2) Using SYNTHEGO Software**

Fragments A–D correspond to the amplified regions containing the target sites of sgRNAs 1, 2, 3, and 4, respectively. Sanger sequencing results show both edited (Ev.2) and wild-type (WT) sequences, with a particular focus on the region surrounding the guide sequence. Mixed bases are frequently observed, indicating the presence of edited alleles with insertions and/or deletions compared to the wild-type allele. The guide sequence is highlighted with a black underline, while the PAM site is marked with a red underline. The expected Cas9 cleavage site is indicated by a vertical black dashed line, where error-prone repair processes often result in the mixed base signals observed after cleavage.

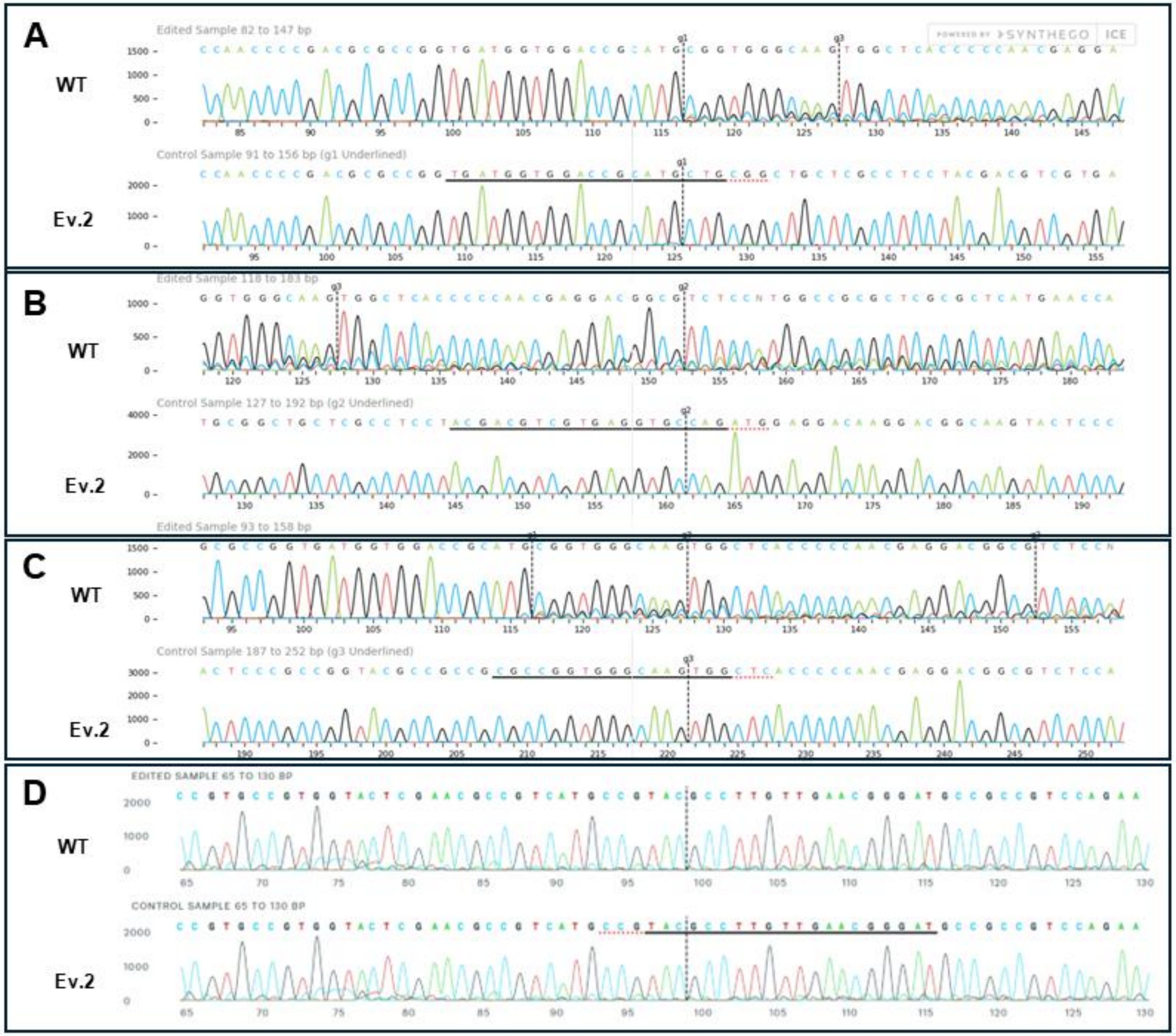


**Supplementary Fig. 4. Sanger Sequencing Chromatogram Analysis of *COMT1* Knockout (Ev.3) Using SYNTHEGO Software**

Fragments A–D correspond to the amplified regions containing the target sites of sgRNAs 1, 2, 3, and 4, respectively. Sanger sequencing results show both edited (Ev.3) and wild-type (WT) sequences, with a particular focus on the region surrounding the guide sequence. Mixed bases are frequently observed, indicating the presence of edited alleles with insertions and/or deletions compared to the wild-type allele. The guide sequence is highlighted with a black underline, while the PAM site is marked with a red underline. The expected Cas9 cleavage site is indicated by a vertical black dashed line, where error-prone repair processes often result in the mixed base signals observed after cleavage.

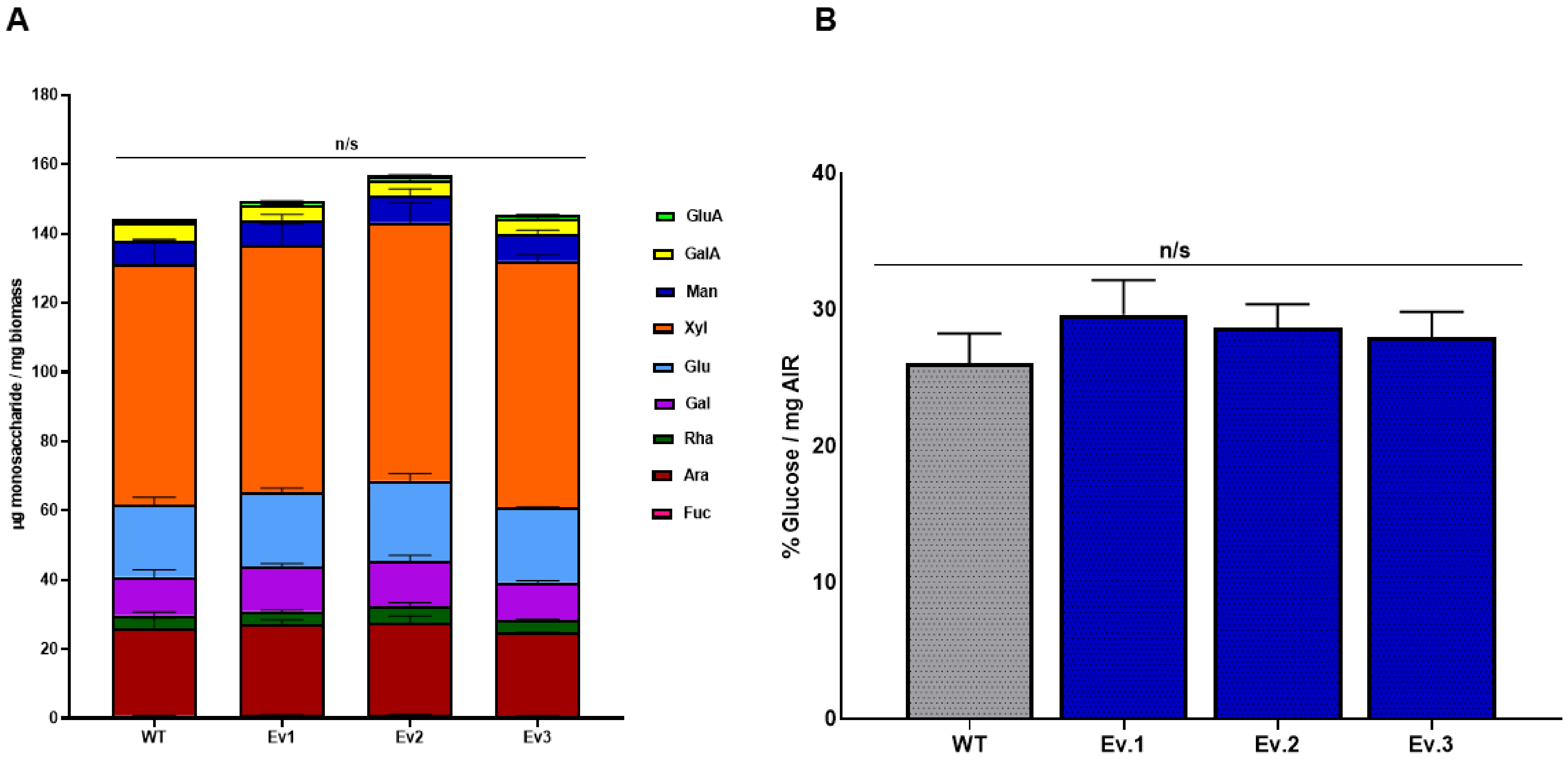


**Supplementary Figure 5. Analysis of monosaccharide composition and cellulose content in wild-type and *COMT1* knockout lines.**

(A) Monosaccharide composition of cell wall polysaccharides, expressed as µg/mg of alcohol-insoluble residue (AIR), and (B) cellulose content, expressed as the percentage of glucose equivalents relative to AIR, in wild-type (WT) plants and *COMT1* knockout lines (Ev.1, Ev.2, and Ev.3). Values are presented as means ± SEM (n = 4). Statistical significance relative to WT was determined by one-way ANOVA followed by Tukey's post hoc test ($p \le 0.05$); n.s., not significant.

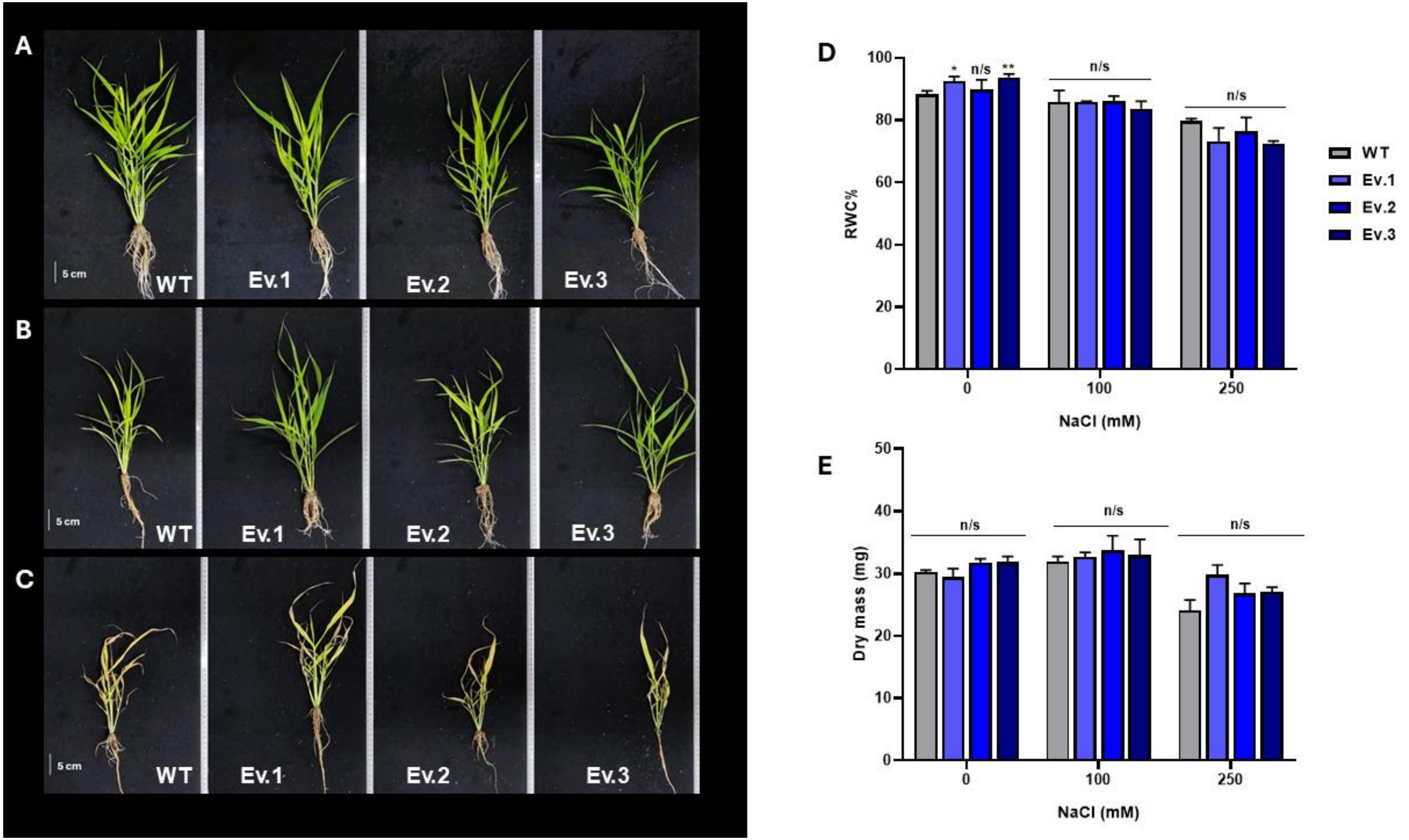


**Supplementary Figure 6. Responses of Relative Water Content and Dry Mass in Wild-Type and *COMT1* Knockout Plants to 21-day NaCl Treatments**

(A–B) Representative phenotypes after 21 days of exposure to 0, 100, and 250 mM NaCl. (C–D) Relative water content (RWC) and dry mass after 21 days under the indicated NaCl concentrations. Values are presented as means ± SEM (n = 4). Statistical significance relative to WT was determined by one-way ANOVA followed by Tukey's post hoc test ($p \leq 0.05$); n.s., not significant.
