## Supplementary material for "Gene Editing of Caffeic-O-methyltransferase (COMT1) in the model grass Setaria viridis Improves Biomass Saccharification Without Compromising Plant Growth or Abiotic Stress Tolerance": Suppl Tables S1-S3

**Supplementary Table 1.** Guide RNA (sgRNA) sequences designed for the knockout of the *COMT1* gene. *Sequences highlighted in bold with an asterisk represent the 3'-5' orientation*

| | ***Sequence*** |
|---|---|
| ***sgRNA1*** | TGATGGTGGACCGCATGCTG |
| ***sgRNA2*** | ***ACGACGTCGTGAGGTGCCAG **** |
| ***sgRNA3*** | ***CGCCGGTGGGCAAGTGG **** |
| ***sgRNA4*** | ATCCCGTTCAACAAGGCGTA |

**Supplementary Table 2.** Primer list used to confirm the transformed events

| **Primer** | **Targuet** | **Sequence** | **Tm (°C)** | **Amplicon size** | **Description** |
|---|---|---|---|---|---|
| **WS28Fw** | cas9 | GAGAAGGGCAAGAGCAAGAA | 62 | 500 | Primer for amplifying the Cas9 gene |
| **WS28Rv** | cas9 | GTGAGGGTGAACAAGTGGATAA | 62 | | |
| **WS29Fw** | hptII | CTGCCTATTCCCGAAGTCCT | 62 | 244 | Primer for amplifying the Cas9 gene |
| **WS29Rv** | hptII | CGCAGATAAAGTCCCTCCAA | 62 | | |
| **WS12Fw** | SvCOMT1 Sevir.6G053300 | GCCGATGACGCTCAAGAA | 62 | 312 | Primer designed to amplify the genomic region flanking sgRNAs 1, 2, and 3 |
| **WS12Rv** | SvCOMT1 Sevir.6G053301 | AGGACCTTGTCCTGGTTCA | 62 | | |
| **WS4Fw** | SvCOMT1 Sevir.6G053302 | CGCTCATGAACCAGGACAA | 62 | 1104 | Primer designed to amplify the genomic region flanking sgRNA 4 |
| **WS4Rv** | SvCOMT1 Sevir.6G053303 | CGAAGCCCGTGTAGAACTC | 62 | | |

***Supplementary Table 3.*** Percentage of indels and knockout score in independent $T_2$ lines

| ***Event $T_2$*** | ***sgRNA 1, 2 and 3 (%indel)*** | ***sgRNA 1, 2 and 3 (knockout score)*** | ***$R^2$*** | ***sgRNA 4 (%indel)*** | ***sgRNA 4 (knockout score)*** | ***$R^2$*** |
|---|---|---|---|---|---|---|
| ***Ev.1*** | 91 | 91 | 0.97 | 0 | 0 | 1 |
| ***Ev.2*** | 75 | 75 | 0.97 | 0 | 0 | 0,99 |
| ***Ev.3*** | 88 | 88 | 0.99 | 0 | 0 | 1 |

The **indel percentage** represents the fraction of sequences with insertions and/or deletions relative to the wild-type sequence. The **Knockout Score** estimates the proportion of alleles with mutations that are likely to result in loss of function of the *COMT*1 gene.
